# α-Synuclein facilitates clathrin assembly in synaptic vesicle endocytosis

**DOI:** 10.1101/2020.04.29.069344

**Authors:** Karina J. Vargas, P. L. Colosi, Eric Girardi, Jae-Min Park, Sreeganga S. Chandra

## Abstract

α-Synuclein and family members β-, and γ-synuclein, are presynaptic proteins that sense and generate membrane curvature, properties important for synaptic vesicle (SV) cycling. αβγ-synuclein triple knockout (KO) neurons exhibit SV endocytosis (SVE) deficits. Here, we investigate how SVE is regulated by α-synuclein. Immuno-electron microscopy (EM) of synaptosomes reveals that α-synuclein relocalizes from SVs to the synaptic membrane upon stimulation, allowing α-synuclein to function on presynaptic membranes during or after stimulation. On cell membranes, we observe that α-synuclein is colocalized with clathrin and its adaptor AP180. Clathrin patches that contain both α-synuclein and AP180 were significantly larger than clathrin patches containing either protein alone. We also find that recruitment of clathrin and AP180 recruitment to membranes are altered in the absence of synucleins. Visualizing clathrin assembly on membranes using an *in vitro* endocytosis reconstitution system reveals that α-synuclein increases clathrin patch size and enhances clathrin lattice curvature, facilitating normal clathrin coated pit maturation. Thus, α-synuclein is an endocytic accessory protein that acts at early stages of SVE to controls the size and curvature of clathrin structures on the membrane.

## Introduction

α-Synuclein became a principal focus of neurodegeneration research when it was identified as the major constituent of Lewy Bodies, pathological protein aggregates found in the brains of Parkinson’s Disease (PD) patients (Spillantini et al., 1997). The importance of α-synuclein to PD pathogenesis was further underscored by the identification of families with Mendelian forms of PD that arise from causal point mutations and gene multiplications of *SNCA*, the α-synuclein gene (Ferese et al., 2015; Ibanez et al., 2004; Kruger et al., 1998; Polymeropoulos, 1997; Singleton et al., 2003; Zarranz et al., 2004). Genome-wide association studies demonstrate that sequence variants in *SNCA* are also associated with sporadic PD (Satake et al., 2009; Simon-Sanchez et al., 2009). Based on these observations, many current therapeutic strategies (Brundin et al., 2017; Tonges & Zella, 2019) for PD are focused on eliminating or reducing α-synuclein levels in the brain. Therefore, there is a growing interest in elucidating the understanding physiological functions of α-synuclein and how loss of α-synuclein impacts neuronal physiology.

Since the discovery of α-synuclein as an SV-associated protein in the electric organ of *Torpedo* (Maroteaux et al., 1988), several distinct approaches have been used to determine its physiological function(s). Structural studies revealed that α-synuclein can adopt several conformations; it is principally unfolded in solution but adopts α-helical structures when associated with phospholipid membranes. In α-helical conformations, the N-terminus folds into either a single elongated amphipathic helix on flatter membranes or a ‘broken’ helix when bound to curved lipid membranes (Chandra et al., 2003; Jao et al., 2008; Lokappa & Ulmer, 2011). α-Synuclein can switch between the two membrane conformations (Ferreon et al., 2009) but has a stronger affinity for highly curved membranes (Middleton & Rhoades, 2010). These biophysical properties are shared by β-, and γ-synuclein and allow these proteins to sense and generate membrane curvature.

The initial description of α-synuclein in *Torpedo* itself suggested an α-synuclein function related to SV trafficking. The membrane-bending properties of synucleins suggest that they regulate SV exocytosis and/or endocytosis. Biochemical studies indicate a role for α-synuclein in exocytosis (Burré et al., 2014; Burre et al., 2010). For instance, α-synuclein has been shown to interact with synaptobrevin-2, a v-SNARE protein required for SV fusion (Burre et al., 2010; Diao et al., 2013; Sun et al., 2019). Somewhat surprisingly, electrophysiological studies suggest that neurotransmission is normal or even improved in αβγ-synuclein KO mice (Anwar et al., 2011; Burre et al., 2010; Greten-Harrison et al., 2010), raising questions about the relevance of this interaction for exocytosis. Functional studies have convincingly demonstrated α-synuclein’s role in modulating SV mobilization and recycling, specifically in SVE (Busch et al., 2014; Medeiros et al., 2018; Schechter et al., 2020; Vargas et al., 2014; Vargas et al., 2017; Xu et al., 2016). Additionally, analysis of SV-like larger diameter neuronal organelles called dense core vesicles in synuclein KOs implicate this family of proteins in fusion pore expansion (Logan et al., 2017). Furthermore, imaging and biochemical analysis have shown striking effects on synaptic clustering and synapsins expression upon deletion of synucleins (Atias et al., 2019; Vargas et al., 2017). Taken together, the preponderance of functional studies supports synucleins acting post-fusion, in SVE, and in SV clustering.

Through pHluorin imaging and cholera toxin labelling, we determined that αβγ-synuclein KO hippocampal neurons exhibit slower SVE kinetics (Vargas et al., 2014). Our unbiased proteomic studies of synaptosomes from αβγ-synuclein KO mice also show significant changes in SVE proteins—increases in clathrin and endophilins and a decrease in levels of the v-SNARE and clathrin adaptor protein AP180 (neuronal homolog of CALM) (Greten-Harrison et al., 2010; Vargas et al., 2014; Westphal & Chandra, 2013). Proteomic analyses of α-synuclein-interacting proteins using proximity labelling in cortical neurons also revealed SVE as one of the most robustly represented pathways, corroborating its involvement in this process (Chung et al., 2017). In addition, we previously demonstrated that the interaction between α-synuclein and synaptobrevin-2 observed by other researchers (Burre et al., 2012; Burre et al., 2010; Sun et al., 2019) is part of a larger complex, of which AP180 is a major component (Vargas et al., 2014). AP180 is a main recruiter of clathrin to synaptic membranes along with AP2, both of which act to initiate clathrin coated pit (CCP) formation. These adaptors are critical for both clathrin assembly and SV protein sorting (Ford et al., 2001; Koo et al., 2015; Koo, Puchkov, et al., 2011; Moshkanbaryans et al., 2014). Specifically, AP180 is responsible for the retrieval of the v-SNARE synaptobrevin-2 to newly formed SVs (Koo, Markovic, et al., 2011). Thus, the transient α-synuclein-AP180-synaptobrevin2 complex may be involved in post-fusion SV protein retrieval. Congruently, AP180 KO mice (Koo, Puchkov, et al., 2011) show synaptic ultrastructural changes similar to αβγ-synuclein KO mice (Greten-Harrison et al., 2010).

Here, we report our investigation of how SVE is regulated by α-synuclein. Our studies reveal that the subsynaptic localization of α-synuclein is dynamic; upon stimulation, α-synuclein relocalizes from SVs to the presynaptic membrane, where it co-localizes with clathrin. We also observed that recruitment of clathrin and its adaptor, AP180, to synaptic membranes is altered in the absence of synucleins. Visualization of clathrin assembly on membranes, using a variety of microscopic methods and an *in vitro* endocytic reconstitution system reveal that α-synuclein functions with AP180 to increase clathrin patch size and curvature, facilitating clathrin-coated pit maturation during SVE.

## Results

### Subsynaptic localization of α-synuclein is dynamic

α-Synuclein is predominantly found on SVs at rest (Boassa et al., 2013; Clayton & George, 1999; Vargas et al., 2017), consistent with its preferential binding to highly curved membranes (Antonny, 2011; Rhoades et al., 2006). To determine whether its localization is altered upon synaptic stimulation, we prepared synaptosomes from wild type mouse brains, and tested three conditions: rest, depolarization with high K^+^ stimulation, and repolarization. Synaptosomes were then immunolabelled with an α-synuclein- or synaptobrevin-2-specific antibody, treated with gold-labelled secondary antibodies and imaged using electron microscopy (EM). At rest, the majority of α-synuclein is present on SVs (61.1% on SVs, 25.9% on synaptic membranes; 5.9% in cytosol; **Fig 1A, B**), similar to the integral SV protein synaptobrevin-2 (78.2% on SV, 12.0% on membranes, 3.1% in cytosol; **Fig 1A, C**), in congruence with previous publications (Vargas et al., 2017). However, upon stimulating synaptosomes with high K^+^, α-synuclein relocates from SVs to the synaptic plasma membrane and cytosol (30.1% on SVs, 51.1% on membranes, 11.2% in the cytosol; α-synuclein on SVs rest versus K^+^ stimulation: 61.1% versus 30.1%; p<0.001). Similarly, previous confocal imaging shows α-synuclein becoming diffuse upon neuronal stimulation (Fortin et al., 2005; Wang et al., 2014). Synaptobrevin-2 remains mainly on SVs when stimulated (rest versus K^+^ stimulation: 78.2% versus 68.4% on SVs; p= 0.095). Repolarization results in α-synuclein relocalizing to the surface of SVs (70.1% on SVs, 18.2% on membranes, 3.8% in cytosol; K^+^ versus recovery: 30.1% versus 70.1% on SVs; p<0.001). To validate that the susbsynaptic localization of α-synuclein is dynamic, we measured the distance from the gold particle to the nearest SV. The distances for α-synuclein at rest, high K^+^, and repolarization were 7.73 nm, 11.32 nm and 4.61 nm, respectively. Gold particle distances for synaptobrevin-2 did not exhibit significant changes and were 10.22 nm, 9.47 nm, and 8.8 nm, respectively. Thus, the sub-synaptic location of α-synuclein is dynamic and is likely to be linked to the SV cycle.

**Figure 1.**
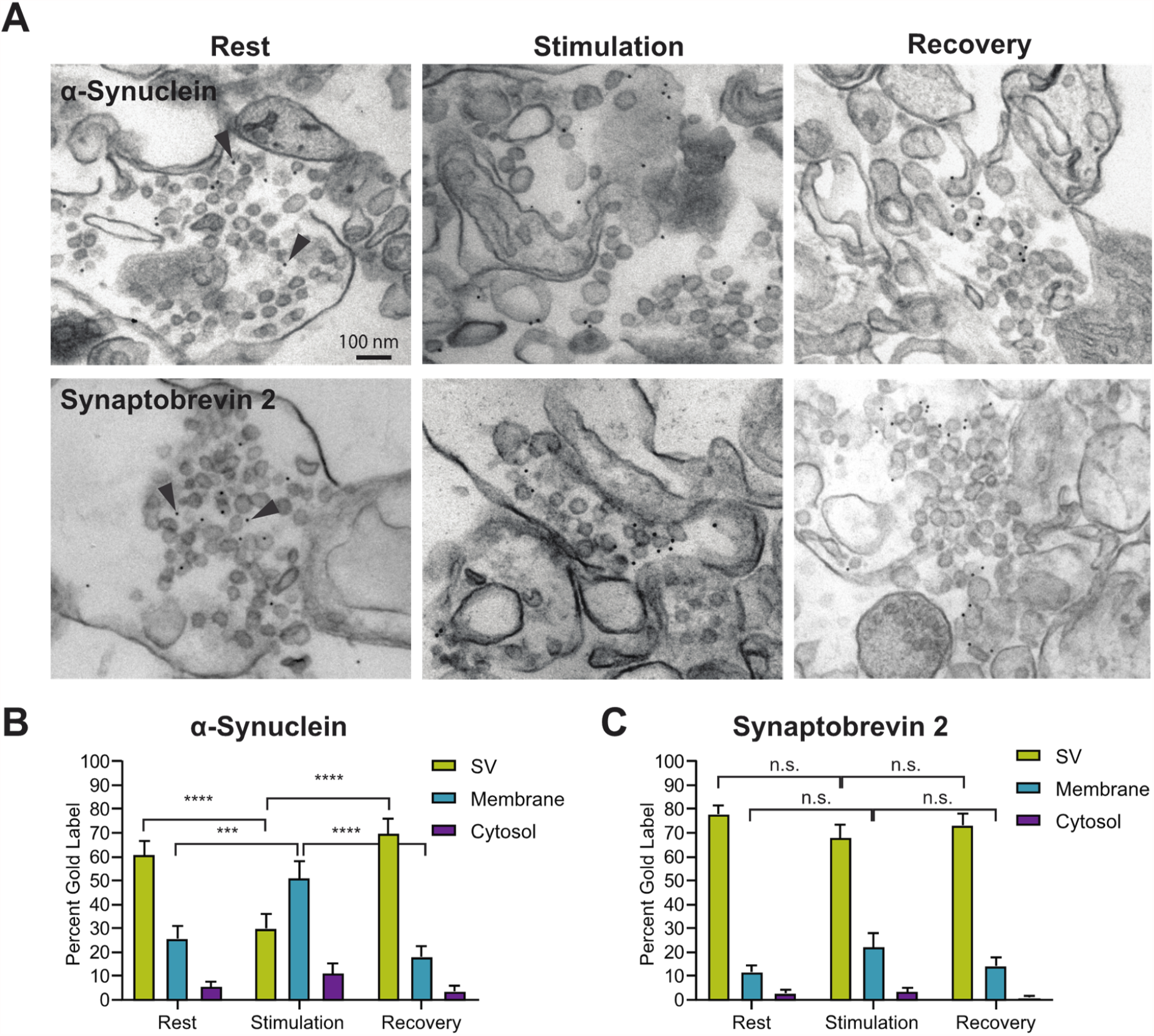
α-Synuclein localization is dynamically regulated by neuronal activity. (**A**) Electron micrographs of permeabilized synaptosomes at rest, stimulated with 90 mM KCl, and upon recovery. Immuno-gold labeled α-synuclein is localized to SVs at rest (arrowhead). During stimulation, α-synuclein disassociates from SVs and predominantly localizes to the synaptic membrane. During recovery, α-synuclein returns to the SV membrane. (**B**) Electron micrographs of permeabilized synaptosomes during rest, stimulation, and recovery conditions immuno-labeled for synaptobrevin-2. Gold-labeled synaptobrevin-2 remains associated with the vesicle in all three conditions. (**C-D**) Quantification of gold label localization for α-synuclein and synaptobrevin-2. Bars represent percent gold label from n=3 independent experiments for control conditions and n=2 for remaining conditions. 25-40 micrographs per condition were analyzed. ANOVA was used to determine significance. * p<0.05; ** p<0.01 and *** p<0.0001. Scale bar=100 nm.

α-Synuclein localization to the synaptic plasma membrane upon neuronal stimulation supports the notion that α-synuclein functions during SV endocytosis. To probe this possibility, we examined α-synuclein location in dynamin 1, 3 double knockout (KO) synapses, which feature few SVs and numerous clathrin coated pit (CCP)-decorated invaginations of the synaptic plasma membrane. The abundance of these invaginations is due to a deficit in scission of CCPs, a process that is needed to form clathrin coated vesicles and ultimately SVs. This membrane trafficking deficit leads to the capture of SVE proteins in CCPs, visualized as clustered in confocal images, while exocytic and peripheral SV proteins, such as rab3a and synapsin, appear unchanged or diffuse, respectively (Ferguson et al., 2007; Raimondi et al., 2011). α-Synuclein appears clustered and co-localizes with clathrin and other endocytic proteins (**Fig. S1**), consistent with our previous findings (Vargas et al., 2014). This suggests that α-synuclein is an accessory endocytic protein that functions prior to dynamin recruitment.

### α-Synuclein and clathrin are co-localized on membranes

To visualize α-synuclein co-localization with clathrin, we prepared adherent membranes by unroofing PTK2 (*Potorous tridactylis* epithelial kidney) cells grown on coverslips transfected with a construct expressing membrane tethered-GFP. We then incubated the adherent membranes with cytosol isolated from WT mouse brain, GTPγS (to prevent dynamin function), and an ATP generation system (t=0-30 minutes) (Wu et al., 2010). We immunostained for clathrin and α-synuclein, then visualized the membranes by light microscopy. At t=0, clathrin and α-synuclein are present in puncta on the membrane (**Fig. 2A-C**), similar to observations in other cell types (Kaur & Lee, 2020; Saffarian et al., 2009). A fraction of puncta contains both proteins. All such puncta display a distinct pattern of a clathrin patch with α-synuclein colocalized at its inner core (**Fig. 2B**). We measured fluorescence intensity and size of clathrin and α-synuclein patches for each punctum. We plotted the relative clathrin and α-synuclein fluorescence for all puncta (n=2071) and binned them into five size categories based on the diameter of clathrin patches (**Fig. 2D**). These data were plotted as a function of clathrin diameter (**Fig. 2E**). For a given clathrin puncta size, the average clathrin fluorescence exceeded that of α-synuclein, suggesting that α-synuclein is found within clathrin patches. Average clathrin fluorescence increases linearly with puncta size, as expected. However, average α-synuclein fluorescence is largely constant and only increases in very large patches (diameter > 1.6 μm) (**Fig. 2E**, blue and purple). This imaging data shows α-synuclein colocalizes with clathrin, predominantly in large puncta.

**Figure 2.**
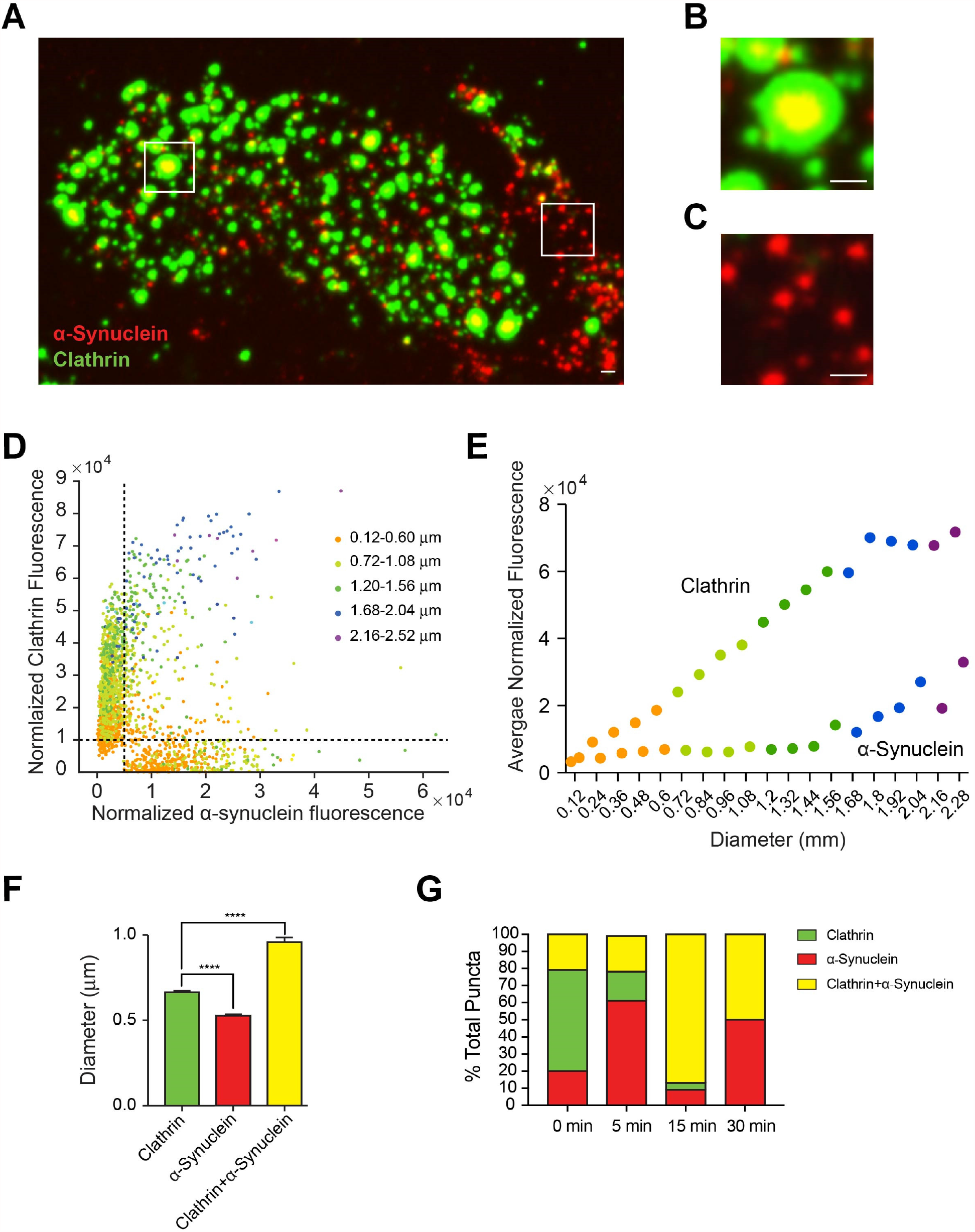
α-Synuclein and clathrin co-localize in patches on the plasma membrane. (**A**) Membrane sheets from PTK2 cells immunostained for α-synuclein (red) and clathrin heavy chain (green). α-Synuclein and clathrin are present in patches on the membrane, with some patches containing both proteins. Scale bar= 1 μm. (**B**) Enlarged view of a clathrin+α-synuclein patch. Note that α-synuclein localizes to the center of the clathrin patch. (**C**) Enlarged view of α-Synuclein only patches. (**D**) Plot of fluorescence of clathrin versus α-synuclein colored by size of the clathrin patch. The dashed lines (x=0.5 and y=1) indicate the cut-offs used to demarcate patches into clathrin only (x<0.5, y>1), α-synuclein only (x>0.5, y<1), and clathrin+α-synuclein (x>0.5, y>1) puncta. Note that clathrin and α-synuclein colocalize predominantly in larger size patches. (**E**) Intensity of clathrin and α-synuclein in different patch sizes. (**F**) Average size of clathrin only, α-synuclein only, and clathrin+α-synuclein patches. (**G**) Changes in the three population of patches over the time when incubated with WT purified cytosol. N=3 independent experiments. * p<0.05; ** p<0.01 and *** p<0.0001. Scale bar=1 μm.

To classify the puncta into clathrin only, α-synuclein only, and clathrin+α-synuclein puncta, we set cutoffs at x=0.5 and y=1 (**Fig. 2D**). Using these cutoffs, 20% of puncta were positive for α-synuclein only, 59% for clathrin only, and 21% for both proteins (**Fig. 2G**; t=0). The average diameters for α-synuclein only, clathrin only, and clathrin+α-synuclein puncta were 0.53 μm, 0.66 μm, and 0.96 μm, respectively (**Fig. 2F**; n=1174, p=0.001), indicating that clathrin+α-synuclein puncta are consistently larger than clathrin-only puncta. Next, we tested how the distribution and size of three puncta classes changed over time during incubation with brain cytosol (**Fig. 2F, S2A**). We observed that the fraction of clathrin only puncta decreases (59% at t=0 to 0% at t=30) with a concomitant increase in clathrin+α-synuclein puncta (20% at t=0 to 50% at t=30), indicating that α-synuclein and clathrin might be co-assembling (**Fig. 2G, S2B**). The average size of α-synuclein-only and clathrin-only puncta remained constant, while the average clathrin+α-synuclein puncta diameter changed over time (**Fig. 2F, S2A**; 0.96 μm at t=0 and 0.77 μm at t=30), mirroring the dynamics of clathrin+α-synuclein puncta.

We extended the adherent membrane studies to include AP180 (Koo et al., 2015), as it can transiently interact with α-synuclein (Vargas et al., 2014). We incubated unroofed membranes with mouse brain cytosol for 15 minutes, because we observed maximum colocalization of clathrin+α-synuclein at this time point. (**Fig. 2G**) When we immunostained for clathrin, α-synuclein, and AP180, we observed a heterogenous population of puncta (**Fig. 3A**). As described above, we classified the puncta into those containing individual proteins (clathrin, α-synuclein, AP180) and two or more proteins (**Fig. 3C**). As expected, we see clathrin alone, α-synuclein alone, AP180 alone, clathrin+α-synuclein and clathrin+AP180 puncta were present (**Fig. 3C**). Surprisingly, we did not observe AP180+α-synuclein puncta, suggesting AP180 and α-synuclein do not interact directly. Intriguingly, 71.4% of puncta contained all three proteins, in concentric circles with AP180 marking the center of the clathrin patch (**Fig. 3A**, see inset). AP180 occupied on average 23% of the clathrin puncta, while α-synuclein puncta occupied 49% of clathrin puncta. We confirmed that concentric ring pattern is seen in all puncta which contain the 3 proteins by measuring the mean diameter for each protein. We observe a hierarchy of diameters with clathrin> α-synuclein > AP180 (**Fig. 3B**). The size of the clathrin shell in clathrin+α-synuclein+AP180 puncta were significantly larger (1.07 µm ± 0.1, p < 0.05) than either clathrin+α-synuclein (0.71 µm ± 0.03) or clathrin+AP180 puncta (0.71 µm ± 0.55)(**Fig. 3E**), suggesting that α-synuclein and AP180 act synergistically to increase clathrin patch size. As noted earlier, the puncta with individual proteins are significantly smaller (**Fig. 3E**)

**Figure 3.**
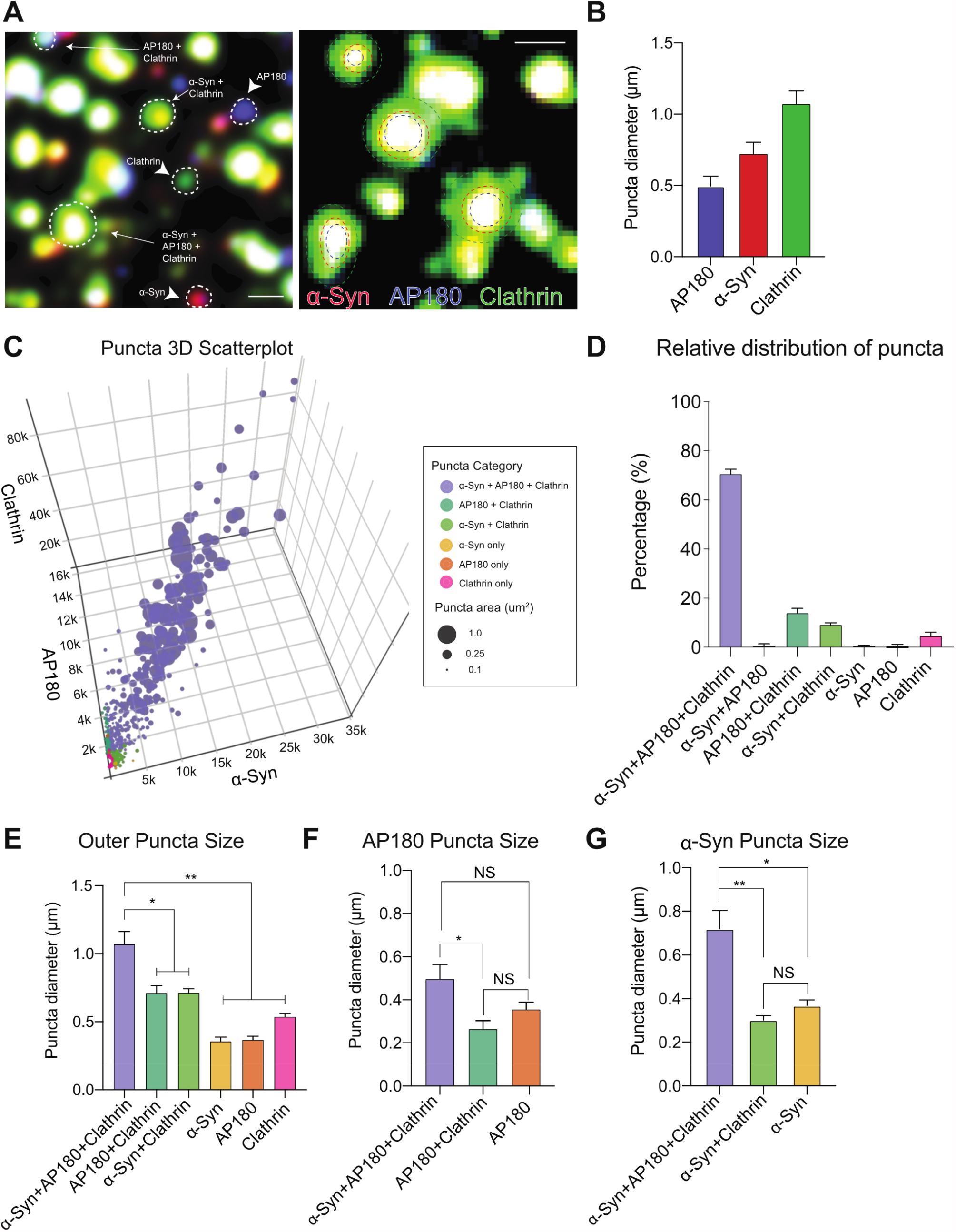
AP180 co-localizes with α-synuclein and clathrin puncta on the plasma membrane. (**A**) Membrane sheets from PTK2 cells immunostained for α-synuclein (red), clathrin heavy chain (green), and AP180 (blue) different classes of puncta are highlighted. Inset showing α-synuclein+AP180+Clathrin puncta. AP180 localizes to the center of the clathrin patch and is surrounded by α-synuclein, denoted by dashed circles. (**B**) Diameter of individual protein puncta in α-synuclein+AP180+Clathrin patches. (**C**) 3D Scatterplot of fluorescence of clathrin versus α-synuclein versus AP180. Individual points are color-coded by puncta identity and their relative size indicates the outer size of puncta. (**D**) Relative distribution of puncta by colocalization identity (**E**) Puncta area by colocalization category (α-synuclein+AP180+Clathrin: 1.07 ± 0.09 µm, AP180+Clathrin : 0.71 ± 0.05 µm, α-synuclein+Clathrin: 0.71 ± 0.03 µm, α-synuclein : 0.37 ± 0.03 µm, AP180 : 0.36 ± 0.06 µm, Clathrin : 0.54 ± 0.03 µm) (**F**) Size of AP180 puncta in clathrin patches containing and lacking α-synuclein. (**G**) Size of α-synuclein puncta in patches containing and lacking AP180. N=3 independent experiments, total number of puncta analyzed = 1681. * p<0.05. ** p<0.01; Scale bar=1 μm.

Next, we monitored AP180 and α-synuclein sizes within the various classes of puncta. The diameter of the AP180 core was larger in clathrin+α-synuclein+AP180 (0.50 µm ± 0.07, p < 0.05) puncta than clathrin+AP180 (0.26 µm ± 0.04) (**Fig. 3F**). The same pattern was observed for α-synuclein (0.72 µm ± 0.08 vs. 0.30 ± 0.02, p < 0.05), strongly suggesting that AP180 and α-synuclein cooperate to increase each other’s patch size within clathrin puncta (**Fig. 3G**). This imaging data suggests that AP180 and α-synuclein function in conjunction to induce larger clathrin patches.

To further test clathrin and α-synuclein co-localization on synaptic membranes, we purified synaptosomes from the brains of WT and αβγ-synuclein KO mice. We identified proteins in proximity with α-synuclein by treating the synaptosomes with a membrane permeable crosslinker, and immunoprecipitating the crosslinked proteins with an α-synuclein antibody (**Fig. S3**). Unique bands of immunoprecipitated proteins from WT samples were processed for mass spectroscopy. The αβγ-synuclein KO samples served as negative controls to identify non-specific interactions. This unbiased analysis revealed that clathrin heavy chain was present in WT samples but absent in the αβγ-synuclein KO samples (**Fig. S3**), indicating that events observed on PTK2 membranes might also occur on synaptic membranes.

### α-Synuclein and clathrin recruitment to membranes

To interrogate α-synuclein involvement in endocytic protein recruitment to the synaptic membrane, we performed an *in vitro* protein recruitment assay. Synaptic membranes isolated from WT mouse brain were stripped of peripheral proteins and incubated with cytosol obtained from WT or αβγ-synuclein KO mouse brains. αβγ-synuclein KO mouse brains show compensatory increases in clathrin, endophilin, and synapsins, decreases in AP180 and the short isoform of AAK1 (**Fig. 4**; (Vargas et al., 2014)). The incubations were supplemented with an ATP generation system and GTPγS, as for PTK2 membrane experiments (**Figs. 2, 3**). Synapsin, which exhibits increased levels in αβγ-synuclein KO and is known to bind synaptic membranes (Hosaka et al., 1999), was used as a positive control (**Fig. 4**). Synapsin binds membranes in proportion to its expression (1.7 ± 0.1; p=0.017). Similarly, we found increased recruitment of clathrin heavy and light chains when synucleins were absent (1.6 ± 0.2 fold, p=0.0061 for clathrin heavy chain; 3.4 ± 0.4 fold, p=0.0016 for clathrin light chain; **Fig. 4**), reflecting the increased clathrin levels in the αβγ-synuclein KO cytosol (Chandra et al., 2005; Greten-Harrison et al., 2010). Despite decreased AP180 levels in αβγ-synuclein KO cytosol (Vargas et al., 2014), we observed a significant increase of AP180 recruitment to the membrane (1.4 ± 0.1 fold, p=0.033). We tested other key endocytic proteins (AP-2, FBP17, epsin1, Eps-15, AAK1, endophilin-A1, necap, and dynamin 1) in the membrane recruitment assay with no significant results. Thus, synucleins may modulate recruitment of clathrin and AP180, proteins involved in the initiation of SVE, to allow for efficient SV exo- and endocytic coupling.

**Figure 4.**
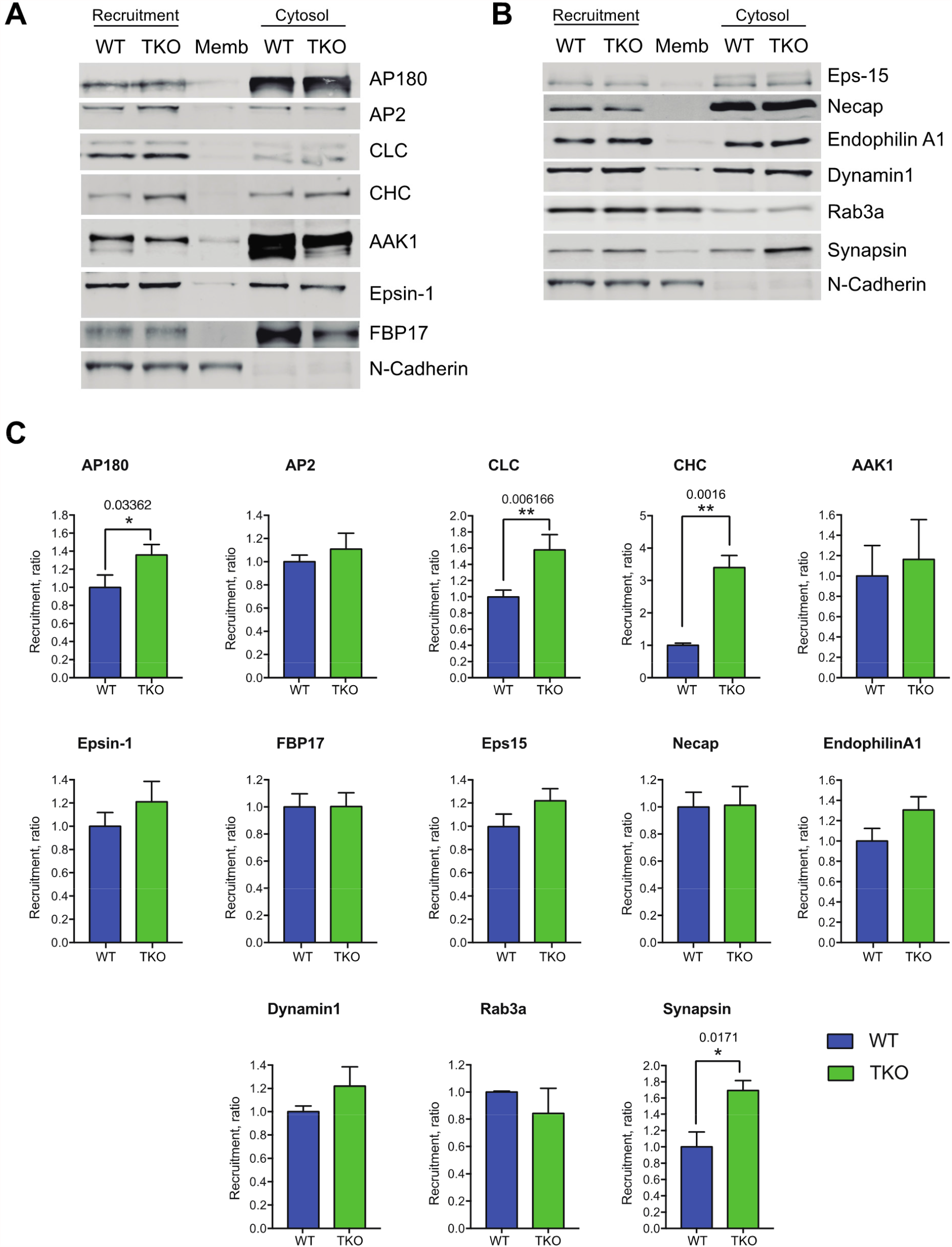
Recruitment of endocytic proteins to synaptic membranes from WT and αβγ-synuclein KO brain cytosol. (**A-B**) Western blot of endocytic and synaptic proteins recruited from WT and αβγ-synuclein KO cytosol to synaptic membranes (Recruitment). Membrane fraction without addition of cytosol is used as a control. N-cadherin was used as a membrane loading control. Abbreviations: Memb, synaptic membrane fraction; CLC, clathrin light chain; CHC, clathrin heavy chain; AAK1, AP2 Associated Kinase-1. (**C**) Quantification of membrane recruitment of endocytic proteins. Western blot quantifications (n=3 independent experiments) after normalizing for added membranes (N-Cadherin signal). * p<0.05, ** p<0.01, Student’s t-test.

To unequivocally establish α-synuclein’s role in clathrin assembly on the membrane, we performed a lipid monolayer assay using purified endocytic proteins (**Fig. 5A**). Addition of clathrin only has no effect on membranes due to its inability to directly bind membranes (**Fig. 5B**), while AP180 can bind the lipid monolayer and modestly deform it (**Fig. 5B**). Injection of recombinant mouse α-synuclein only leads to membrane ruffles and deformations (**Fig. 5C**), consistent with its ability to bend membranes (Westphal & Chandra, 2013). Addition of both AP180 and clathrin leads to the formation of discrete clathrin cages (**Fig. 5D**), confirming previous literature showing AP180 as necessary and sufficient for CCP formation (Ford et al., 2001). Addition of α-synuclein leads to significantly larger clathrin assemblies (**Fig. 5D-F**) (Clathrin+AP180 = 8607 ± 413 nm^2^, Clathrin+AP180+α-synuclein= 15,569 ± 2131 nm^2^; p<0.01), corresponding with the larger patches seen in the PTK2 membrane assay (**Fig. 3A, E**).

**Figure 5.**
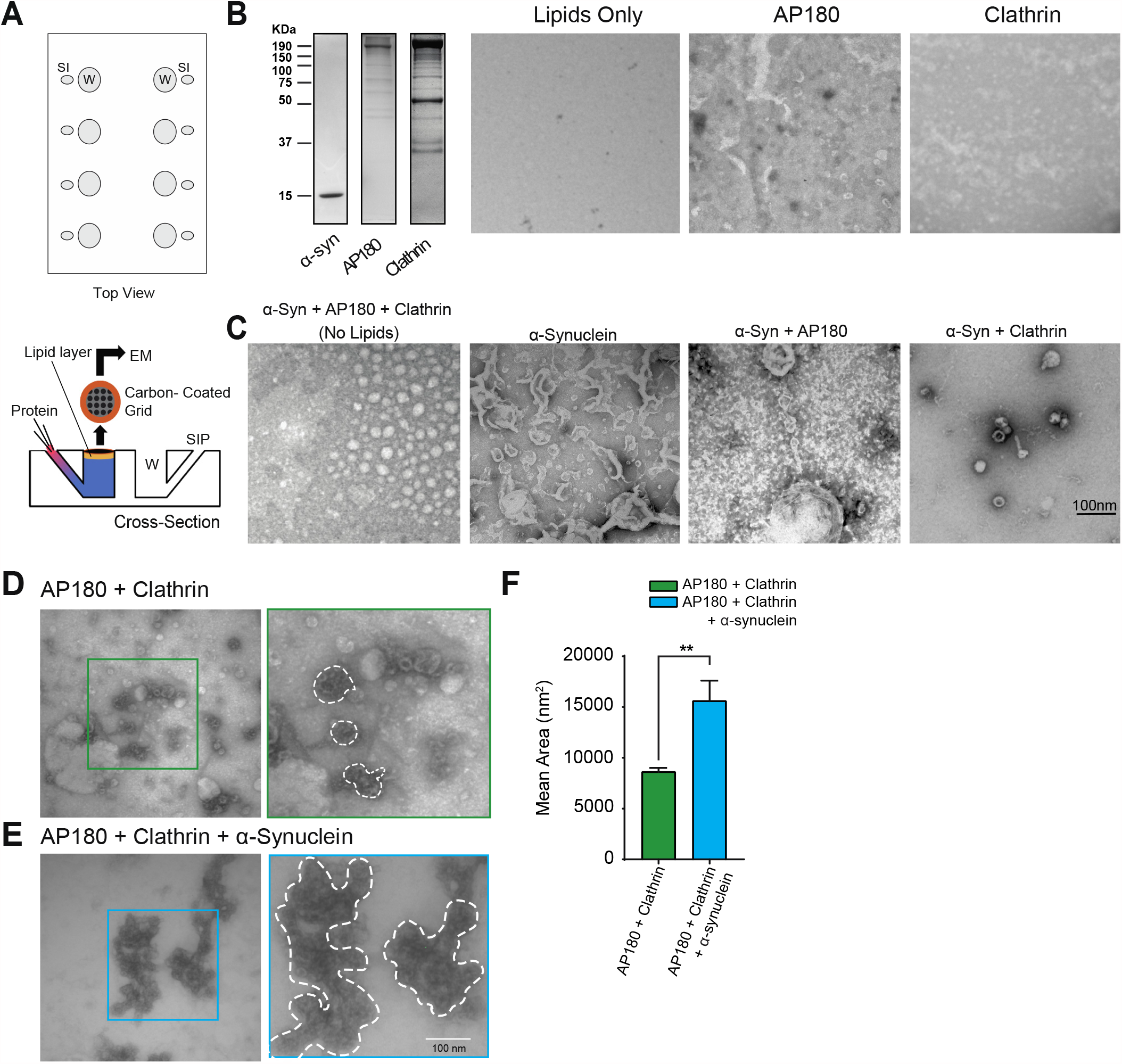
Reconstitution of clathrin assembly using recombinant proteins. (**A**) Schematic of Teflon block and experimental setup used in monolayer reconstitutions. Wells (W) and Side Injection Ports (SIP) labeled. (**B**) From left to right: Coomassie stain showing proteins used in reconstitution conditions and electron micrographs of negative controls; Lipids only: Lipid monolayer with no added protein; AP180: recombinant AP180 added to lipid monolayer; Clathrin: Brain purified clathrin added to lipid monolayer. (**C-E**) Electron micrographs of constituent proteins added to lipid monolayers. **(C)** First panel: All proteins without a lipid monolayer; Other panels; α-Synuclein: Recombinant WT mouse α-synuclein only membrane ruffles are observed; α-Syn+AP180: Recombinant mouse AP180 and α-synuclein; α-Syn+Clathrin: α-Synuclein and isolated clathrin added to lipid monolayer. (**D**) Electron micrographs of clathrin and AP180 added to lipid monolayer. Characteristic fullerene-shaped clathrin cages form. Inset: Higher magnification image outlined in green (**E**) Electron micrographs of α-synuclein, clathrin, and AP180 added to lipid monolayer. Inset: Zoomed image outlined in blue. Scale bar = 100 nm. (**F**) Quantification of area of clathrin patches obtained in conditions **D** and **E**. Addition of α-synuclein increases the area of clathrin patches. N=3 independent experiments with at least 10 micrographs per experiment. * p<0.05; ** p<0.01 and *** p<0.0001. Scale bar=100 nm.

### Examining the role of α-synuclein in clathrin lattice curvature

Our previous SVE kinetic measurements demonstrated slower endocytosis in αβγ-synuclein KO neurons (Vargas et al., 2014). Here, we observed increased membrane recruitment of clathrin and AP180 biochemically *in vitro* (**Fig. 4**). Prior studies have suggested that the clathrin:adaptor ratio is a determinant of CCP curvature (Bucher et al., 2018). The clathrin:AP180 protein ratio on synaptic membranes for αβγ-synuclein KO displayed an 1.14 and 2.42 fold increase for heavy and light chain, respectively, suggestive of altered curvature of assembled clathrin lattices. To reconciliate these findings, we sought to determine the state of clathrin assembly on the membrane in the αβγ-synuclein KO condition.

To visualize clathrin lattice assembly, the lipid monolayer assay designed for incubation with purified proteins (**Fig. 5A**) was adapted for brain cytosol (Ford et al., 2001) (**Fig. 6A**). WT and αβγ-synuclein KO cytosol was used to quantify the time dependence of clathrin assembly into flat lattices and mature CCPs. We observed that incubation of the lipid monolayer with WT cytosol leads to assembly of flat and curved clathrin lattices by 15 minutes (**Fig. 6A, B**), in line with previous publications (Ford et al., 2001). On occasion, mature CCPs were also seen (**Fig. 6A**). The size of individual clathrin patches was quantified in **Fig. 6B**. The absence of synucleins initially leads to significantly smaller clathrin patches (**Fig. 6B**, t=15, WT= 30,808 ± 2,617 nm^2^ and αβγ-synuclein KO = 25,544 ± 2,134 nm^2^, p<0.001). However, by t=23 minutes, the patches are significantly larger (t=23, WT= 36,513 ± 2597 nm^2^, and TKO= 64,882 ± 3790 nm^2^, p<0.05), in line with the increased membrane recruitment of clathrin and AP180 (**Fig. 4**). The delayed clathrin assembly in the αβγ-synuclein KO is consistent with slower SVE described in neurons (Vargas et al., 2014).

**Figure 6.**
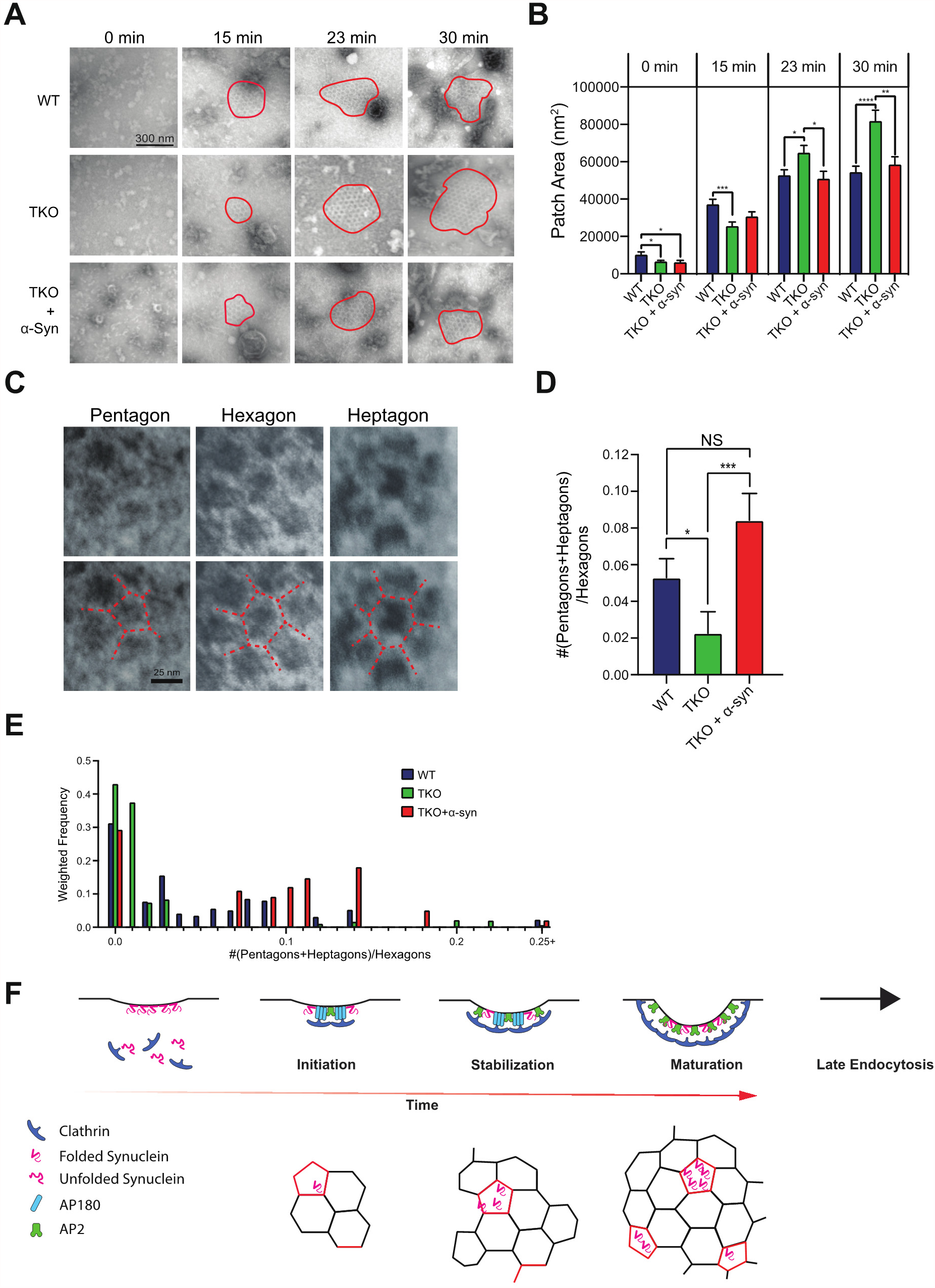
Lipid monolayer assay to visualize clathrin recruitment from brain cytosol. (**A**) Electron micrographs of clathrin assembly into flat lattices and CCPs on membrane monolayers over time (0,15, 23, and 30 min). Membrane monolayers were incubated with cytosol from WT, αβγ-synuclein KO, and mouse α-synuclein added to TKO cytosol (rescue). Clathrin patches are highlighted with red outlines. Arrow indicates CCP. N=3 independent experiments with 10 images per experiment. Scale bar=100 nm. (**B**) Comparison of clathrin patch areas in the three conditions mentioned above. (**C**) Examples of pentagon, hexagon, and heptagon clathrin lattices. (**D**) Ratio of non-hexagons (pentagons and heptagons) to hexagon in clathrin lattices for each of the three conditions, t=23 min. (**E**) Histogram of non-hexagon to hexagon ratio for the three conditions. The sum of all weights was used to normalize the data. For all graphs: * p<0.05; ** p<0.01 and *** p<0.0001. Welsh’s t-test. **(F)** Model of α-synuclein function in clathrin assembly.

We examined the curvature of clathrin patches by analyzing the fractions of pentagons and heptagons to hexagons in the clathrin formations (**Fig. 6C, D**). The presence of pentagons and/or heptagons indicates that a curved lattice is being generated. We found that incubation with αβγ-synuclein KO cytosol leads to fewer non-hexagonal clathrin formations (**Fig. 6D, E**), indicating flatter clathrin lattices. To confirm that clathrin assembly deficits are caused by the absence of synuclein, we performed rescue experiments, by adding recombinant mouse α-synuclein (at endogenous concentrations) to αβγ-synuclein KO cytosol restored both the kinetics of formation and size of clathrin patches, at later time points, to that of WT sizes (t=30, WT= 54,434 ± 3,165 nm^2^, TKO= 81,753 ± 5,840 nm^2^, and TKO + α-synuclein= 58,510 ± 4,127 nm^2^; WT versus TKO + α-synuclein, ns) (**Fig. 6A, B**). In addition, a greater number of lattices with pentagons and heptagons were present in the TKO + α-synuclein condition, shapes which are associated with curved lattices (**Fig. 6D, E**). Together, these results suggest that synucleins function to enlarge and curve the clathrin lattice.

## Discussion

Our findings clearly categorize synucleins as endocytic accessory proteins. Synucleins, similar to other SVE proteins, bind acidic lipids, especially Phosphatidylinositol 4,5-bisphosphate (PI(4,5)P_2_), which is essential for SVE. Our data indicate that the subcellular localization of α-synuclein is dynamic, as it moves to the synaptic membrane upon stimulation (**Fig.1**), where it can participate in SVE. Moreover, in dynamin 1,3 double KO neurons, α-synuclein is clustered with other endocytic proteins (**Fig. S2**). Unbiased proximity labelling (Chung et al., 2017) and crosslinking experiments (**Fig. S3**) suggest that clathrin and other SVE proteins are α-synuclein binding partners on the membrane (Chung et al., 2017), where α-synuclein multimerizes forming puncta (**Fig. 2C**). We suggest that the interactions of α-synuclein are of low affinity and multivalent, like interactions of other SVE proteins (Evergren et al., 2004; Pechstein et al., 2010; Ringstad et al., 1999). Previous work, including from our laboratory, demonstrates that synucleins can effectively sense and generate membrane curvature (Westphal & Chandra, 2013). Functionally, the absence of synucleins leads to slowed SVE (Schechter et al., 2020; Vargas et al., 2014) and enhanced frequency depression due to decreased SV pool recovery (Vargas et al., 2014). In this study, we establish that α-synuclein affects clathrin assembly and curvature generation, pinpointing the steps of SVE in which α-synuclein participates.

How does α-synuclein facilitate CCP formation (**Fig. 6F**)? We propose that α-synuclein and AP180 function synergistically on the membrane (**Fig. 3A, D**), via their interactions with PI(4,5)P2 (Burre et al., 2012; Burre et al., 2010; Sun et al., 2019; Vargas et al., 2014), and α-synuclein’s ability to self-assemble (**Fig. 2C**) and bend the membrane (**Fig. 5C, F**) thereby promoting clathrin patch initiation. This is supported by our discovery of a transient α-synuclein/AP180 complex assembled post-stimulation (Vargas et al., 2014). Our direct observation of α-synuclein surrounding AP180 within larger clathrin patches on cell membranes (**Fig. 2B; Fig 3A**) is also in line with this hypothesis. Once initiated, we speculate that the geometry of α-synuclein and AP180-containing puncta stabilizes the clathrin lattice, allowing for the formation of larger clathrin puncta (**Fig. 5D-E**). This idea is supported by our findings that AP180 and α-synuclein cores within clathrin patches are larger when both proteins are present (**Fig. 3F, G**). Interpreting the results from the αβγ-synuclein KO experiments is complicated by the compensatory changes in the cytosolic levels of clathrin and AP180 (**Fig. 4A, C**) and phosphorylation changes in endocytic proteins (Vargas et al., 2014). Yet, the time course of clathrin lattice formation is initially slower in the αβγ-synuclein KO condition (**Fig. 6B-C**), mirroring the slow kinetics of SVE (Vargas et al., 2014). The larger patches seen in the lipid monolayer assay with the αβγ-synuclein KO cytosol at later time points (**Fig. 6C**) likely occur due to the upregulated levels of clathrin in these mice (**Fig. 4A, C**). We suggest that the larger clathrin patches take longer to fully curve and are shallower. Congruently, we find CCPs with fewer pentagons and heptagons in the αβγ-synuclein KO condition (**Fig. 6D**), indicating shallower CCPs. In the rescue condition, the clathrin patch size is restored (**Fig. 6B**) despite more clathrin in the cytosol, because the resulting patches are more curved (**Fig. 6D-E**). Thus, α-synuclein coordinates clathrin assembly with membrane curvature generation (**Fig. 6F**).

One striking finding is that α-synuclein localizes within clathrin puncta (**Fig. 2**) and is arranged in concentric rings with AP180 at the center (**Fig. 3**). The localization of endocytic proteins in clathrin assembled structures on HeLa adherent membranes is known (Sochacki & Taraska, 2017); Correlative super resolution microscopy combined with EM (CLEM), reveals that most endocytic proteins are localized to the edge of clathrin assemblies on the membrane (both flat lattices and CCPs). The only two proteins found in the middle of the clathrin structures are CALM and VAMP, the non-neuronal homologs of AP180 and synaptobrevin-2 (Sochacki et al., 2017). Our findings suggest that α-synuclein corrals AP180 to the center of clathrin patch on synaptic membranes. This could serve to achieve the high number of synaptobrevin-2 molecules present on SVs (∼70/SV) (Takamori et al., 2006). Supporting this idea, SVs purified from αβγ-synuclein KO brains show lower levels of synaptobrevin-2 compared to WT brains (Vargas et al., 2014). Similarly, in AP180 KO mice, activity-dependent defects in synaptobrevin-2 recycling result in reduced synaptobrevin-2 levels on SVs (Koo et al., 2015). Overall, our results support that α-synuclein regulates CME for proper SV cycling.

## Materials and Methods

### Immuno-electron microscopy of mouse brain synaptosomes

#### Synaptosome Isolation

Brains from WT mice (n=3) were harvested and washed in homogenization buffer (0.32 M Sucrose, 10 mM Tris-HCl pH 7.4, mini cOmplete protease inhibitor). Cerebella were removed, hemispheres separated, and white matter removed before adding the remaining brain to 10 mL of homogenizing buffer. Brains were homogenized in a 55 mL Potter homogenizer at 80% power for 8 strokes. Homogenate was centrifuged at 2,330 x g for 4 min at 4°C in a JA-20 rotor. Pellet (P1) was discarded, and supernatant (S1) was transferred to a 50 mL centrifuge tube. S1 was recentrifuged at 18,850 x g for 12 min at 4°C. Supernatant (S2) was discarded and pellet (P2) was resuspended in 6 mL of homogenization buffer. Gradients of Percoll solutions (from bottom to top: 2.5 mL 23%, 3 mL 10%, and 2.5 mL 3%) were prepared in 15 mL glass tubes. 2 mL of resuspended P2 was added to the top of each gradient, and tubes were centrifuged at 18,850 x g for 12 min at 4°C. Synaptosome fractions were collected by Pasteur pipette.

#### Stimulation

Washing buffer Krebs 1 (140 mM NaCl, 10 mM Tris-HCl pH 7.4, 5 mM KCl, 5 mM NaHCO_3_, 1.3 mM MgSO_4_, 1 mM Na-phosphate buffer) was added to the collected synaptosomes, and sample was centrifuged at 18,850 x g for 12 min. Pellet was resuspended in 1.5 mL Krebs 1 and divided equally into three 1.5 mL microcentrifuge tubes labeled for rest, stimulation, and recovery. The tubes were centrifuged at 16,000 x g for 3 min. Each pellet was resuspended in 500 μL of washing buffer Krebs 2 (10 mM glucose, 1.2 mM CaCl_2_ in Krebs 1) and incubated in a 37°C water bath for 15 min. The rest condition was left in the bath while the stimulation and recovery conditions were each removed, supplemented to 90 mM KCl final concentration, rotated by hand, and returned to the bath for 2 min. Stimulation and recovery tubes were then spun at 16,000 x g for 2 min. The stimulation pellet was resuspended in 200 μL Krebs 1 and left on ice. The recovery pellet was resuspended in 500 μL of washing buffer Krebs 3 (10 mM Glucose, 1 mM EGTA, 1 phosSTOP tablet in Krebs 1) and returned to the bath for 10 min. The recovery and rest tubes were removed from the bath and spun at 16,000 x g for 2 min. Pellets were resuspended in 200 μL of Krebs 1. All conditions were transferred into separate 50 mL centrifuge tubes filled with 30 mL of hypotonic fixative solution (3% paraformaldehyde, 0.25% glutaraldehyde in 5 mM Na-phosphate buffer) and incubated on ice for 30 min.

#### Agarose Embedding

The samples were centrifuged in their 50 mL tubes at 18,850 x g at 4°C for 12 min. Each pellet was resuspended in 400 μL of 120 mM Na-phosphate buffer. 180 μL aliquots of each sample were pipetted into glass test tubes and held on ice until mixed with agarose. Pasteur pipettes and glass slide frames (see (De Camilli et al., 1983)) were warmed in a 60°C oven. 3% Agarose solution was prepared in 5 mM Na-phosphate buffer, warmed to 95°C until agarose dissolved, and kept in a 54°C water bath. 180 μL of agarose solution was pipetted over each broken synaptosome sample while the tube was held in the 54°C water bath and mixed by gentle pipetting with a Pasteur pipette. The agarose-embedded samples were pipetted into the warmed glass slide frames and solidified on ice. Once solid, the frames were disassembled, leaving a solid agarose gel sheet on one glass slide. The gel was cut into 3 mm^3^ cubes with a razor blade and washed from the slide with 120 mM Na-phosphate buffer into petri dishes.

#### Immunolabeling and Electron Microscopy Preparation

Agarose cubes from each condition were divided into 5 wells of glazed porcelain well plates. Cubes were incubated for 30 min with 5% bovine serum albumin (BSA) in solution A (0.5 M NaCl, 0.02 M Na-phosphate buffer) at room temperature. BSA solution was removed, and each well was incubated overnight with 200 μL of primary antibodies in solution A at 4°C. Cubes were washed with 5 changes of solution A over the course of 1.5 hours. Solution A was removed from the wells and cubes were incubated with secondary antibodies in solution A and 5% BSA for 6 hours at room temperature. Cubes were washed with 5 changes of solution A and incubated at 4°C overnight. Agarose samples were prepared for resin embedding, sectioned, and stained with 2% uranyl acetate and 1% lead citrate before imaging on a FEI Tecnai Biotwin scope at 48k, 80kV.

### Reconstitution of endocytosis in PTK2 Cells

PTK2 cells were transfected to express palmitoylated GFP and cultured for 48 hours in a Poly-D-Lysine coated Mattek dish (P35G-1.5-14-C) to confluence. Membrane sheets were prepared by sonication using 20% output power at a 20 mm distance, with 1-2 brief pulses at the center of the well. The sheets were washed gently with cytosolic buffer and used within 20 minutes after unroofing. The membranes were incubated for 0, 5, 15, or 30 minutes with 2 mg/mL cytosol, 1.5 mM ATP, ATP reconstitution system (16.7mM phosphocreatine, 16.7U/ml creatine phosphokinase), and 150 μM GTPγS. The reaction was stopped by gently washing twice with cytosolic buffer, followed by immediate fixation with 4% PFA for 15 minutes in PBS. Membrane-bound proteins were detected using appropriate primary antibodies: goat anti-alpha synuclein (Everest Biotech #EB11713), mouse anti-AP180 (SantaCruz #LP2D11), rabbit anti-clathrin heavy chain (Cell Signaling # d3c6) and appropriate AlexaFluor conjugated secondary antibodies. Z-stack images were captured using spinning disc confocal microscopy and image analyses were performed using Fiji. Z slices were stacked using the sum slice function to create a single composite image. For each circular structure observed on membranes of interest, circular ROIs were manually drawn and analyzed for size and fluorescence intensity to categorize each puncta according to denoted thresholds. After categorization of each puncta, linear ROIs were manually drawn as diameters for each puncta in different channels to measure the size of each protein puncta. 3D scatterplot for fluorescence was generated by utilizing the *plotly* package in R studio (v. 4.0.2) and statistics for individual analysis of puncta category was done using GraphPad 8.

### Isolation of synaptic membranes

All steps were executed at 4°C. Two WT mouse brains were washed in Buffer A (1 mM MgCl_2_, 1 mM CaCl_2_, 1 mM NaHCO_3_, 0.32 M sucrose) for 30 seconds and transferred to a 55 mL potter homogenizer with 6 mL of Buffer A. The samples were homogenized at 5,000 rpm for 10 strokes. The homogenate was centrifuged at 1,400 x g for 10 min. Supernatant was retained. The pellet was resuspended in 6 mL of Buffer A and centrifuged at 700 x g for 10 min. Both supernatants were combined and centrifuged at 17,500 x g for 15 min. The supernatant was discarded, and the pellet was resuspended in 3 mL Buffer A. The resuspended pellet (1.5 mL) was added to the top of freshly prepared 0.65 M, 0.85 M, 1.00 M, 1.20 M sucrose gradients (1.5 mL each concentration). The gradients were centrifuged at 100,000 x g for 2 hours. Synaptosomes were collected from the 1/1.2 M interface and suspended in an excess of Buffer A. The synaptosomes were centrifuged at 100,000 x g for 20 min. The pellet was resuspended in 4 mL ice-cold deionized water, and HEPES pH 7.4 was added to a final concentration of 7.5mM. The suspension was incubated on ice for 30 min and centrifuged at 100,000 x g for 20 min. The pellet was resuspended in 4 mL of 0.1 M Na_2_CO_3_ to strip peripheral proteins, incubated for 15 min at 37°C, and centrifuged at 100,000 x g for 20 min. Pellet was resuspended in 2 mL cytosolic buffer, centrifuged again at 100,000 x g for 20 min, and resuspended in 2 mL cytosolic buffer. Proteins were quantified using BCA bioassay. Mini cOmplete protease inhibitor cocktail was added, and 100 μl aliquots of purified membrane resuspension were flash frozen in liquid nitrogen and stored at −80°C.

### Cytosol purification

Mouse brains were removed and washed in washing buffer (23 mM Tris-HCl, pH 7.4, 320 mM sucrose) for 30 seconds. Two brains were homogenized at 2,500 rpm in a 5 mL potter homogenizer with 2 mL of homogenization buffer (25mM Tris-HCl, pH 8.0, 500 mM KCl, 250 mM sucrose, 2 mM EGTA, and 1 mM DTT), using 10 strokes at 5,000 rpm. The homogenate was transferred to a 3.5 mL ultracentrifuge tube and centrifuged at 160,000 x g for 2 hours at 4°C. A PD-10 column was equilibrated with 25 mL cytosolic buffer, the supernatant was added to the PD-10 column, and then, eluted with 3.5 mL of cytosolic buffer (25 mM Hepes-NaOH, pH 7.4, 120 mM potassium glutamate, 2.5 mM magnesium acetate, 20 mM KCl, and 5 mM EGTA-NaOH, pH 8.0, filtered and stored at 4°C, with 1 mM DTT added immediately before use). Protein concentration was quantified by BSA assay. Mini cOmplete protease inhibitor cocktail was added, and 100 μl aliquots were flash frozen in liquid nitrogen and stored at −80°C.

### Synaptic membrane protein recruitment

Synaptic membranes (400 μg/mL) were mixed with 0.5 mg/mL WT or αβγ-synuclein KO cytosolic proteins in 500 μl cytosolic buffer (see above, with 1 mM DTT added immediately before use) and supplemented with an ATP regenerating system (1.5 mM ATP, 16.7 mM phosphocreatine, and 16.7 U/mL creatine phosphokinase) and 150 μM GTPγS. A control experiment was prepared with 400 μg/mL synaptic membranes in 500μl of cytosolic buffer alone. Mixtures were pipetted 3 times to mix and incubated at 37°C for 15 min. The samples were immediately centrifuged at 100,000 x g for 30 min at 4°C. Pellets were thoroughly resuspended in 500 μl of cytosolic buffer at 4°C. The resuspension was centrifuged at 100,000 x g for 30 min at 4° resuspended in 45 μl of cytosolic buffer. For each sample, 20 μl aliquots were mixed with 5 μl of 5X loading buffer. WT and αβγ-synuclein knockout cytosol reference samples were prepared by diluting 32 μg of each cytosol to 20 μl with cytosolic buffer and mixing with 5 μl of 5X loading buffer. Samples were boiled for 10 min at 100°C. Levels of various proteins were measured for each condition by quantitative immunoblotting.

### Reconstitution of clathrin recruitment on lipid monolayers

#### Purification of proteins

α-Synuclein and AP180 were recombinantly expressed and purified from BL21 (DE3) *E. Coli* as previously described (Ford et al., 2001; Taguchi et al., 2017; Westphal & Chandra, 2013). Clathrin was purified from porcine or bovine brains by purifying CCVs and disassembling cages as described (Chen et al., 2019). For the described experiments, we used porcine and bovine clathrin interchangeably with no significant difference.

#### Lipid Monolayer Assay with Purified Proteins

An 8-well Teflon block was arranged in a humid chamber and wells were filled with 40-60 μL of HKM buffer (25 mM HEPES pH 7.4, 125 mM potassium acetate, 5mM magnesium acetate, 1mM dithiothreitol). Lipid mixture (10% cholesterol, 40% PE, 40% PC, and 10% PI(4,5)P2 to final concentration of 0.1 mg/mL in a 19:1 mixture of chloroform to methanol) was carefully pipetted onto the buffer in each well. The blocks were incubated in a humid chamber at room temperature for 1 hour to evaporate methanol/chloroform. Carbon-coated copper grids were placed carbon-side down onto each well. Proteins were introduced via side-injection ports beside each well. Final protein concentrations per well were as follows: 2 μM AP180 and α-synuclein, 500 nM purified bovine/porcine clathrin. Grids were incubated for 60 min in a humid chamber at room temperature, then removed from the block and immediately negative stained with 1% uranyl acetate. After drying, grids were imaged on a Phillip CM10 transmission electron microscope at 46k, 80kV). Clathrin lattices were outlined manually, and the areas were quantified in Image J. Non-hexagonal clathrin lattices could not be quantified due to the high contrast needed to visualize these structures in these experiments.

#### Lipid Monolayer Assay with Mouse Brain Cytosol

5 mg/mL WT and αβγ-synuclein KO mouse brain cytosol was added to wells of the Teflon block. Lipid mixture (see above) was pipetted on top of the cytosol. Blocks were incubated in a humid chamber for 1 hour to evaporate methanol/chloroform. Carbon-coated copper grids were placed on top of the wells, and an ATP regenerating system (see above) and 150 μM GTPγS were introduced through a side injection port. Grids were incubated in a humid chamber at 37°C for 0, 5, 15, or 30 min and then removed from the block and immediately negative stained as described above. Grids were imaged on a FEI Tecnai Biotwin scope at 42k, 80kV. Clathrin lattices were outlined manually, the area and number of pentagons, heptagons, and hexagons were quantified in Image J.

### Crosslinking experiments

Synaptosomes were isolated from WT and αβγ-synuclein KO mouse brains as described. Synaptosomes were incubated with SDAD crosslinker (1 mM; Thermo) or DMSO as control and incubated on ice for 45 minutes. The reaction was stopped with 100 μl 1M Tris, pH 8.0 on ice for 15 minutes. The excess crosslinker was removed by centrifugation. The synaptosomes were then UV crosslinked for 1 minute with a Strategene UV light (254 nm). Crosslinked synaptic membranes were solubilized with 1% Triton and used as starting material for α-synuclein immunoprecipitation experiments. The immunoprecipitants were run on SDS-PAGE, silver stained with a mass spectrometry compatible silver stain (Pierce), relevant bands were excised and identified by LC-MS.

### Statistics

All experimental analysis was done blinded to condition. Statistical analysis was performed by GraphPad and R studio. Data is presented as average ± SEM.

## Acknowledgments

This work was supported by NIH (R01 NS064963, R01 NS110354, R01 NS083846), Nina Compagnon Hirshfield Parkinson’s Disease Research Fund and DOD (W81XWH-17-1-0564). The proteomic experiments were supported by the NIDA Neuroproteomic Center (NIH DA018343-11A1). We would like to thank Dr. Pietro De Camilli and his lab members for Dynamin 1,3 double KO pups as well as help in performing the PTK2 cell assays. We thank Aurelie Nardin and Jeffery Yong for setting up and optimizing the PTK2 assay, Christopher Westphal and Becket Greten-Harrison for help with cross-linking experiments. We thank John. E. Lee and Sofia Massaro Tieze for editing the manuscript.

## Author contributions

KJV, PC and SSC designed research/experiments; KJV, PC, EG, JP, and SSC performed experiments; KJV, PC, EG, JP, and SSC analyzed data; All authors wrote and edited the paper.

## SUPPLEMENTARY FIGURES

**Supplementary Figure 1.**
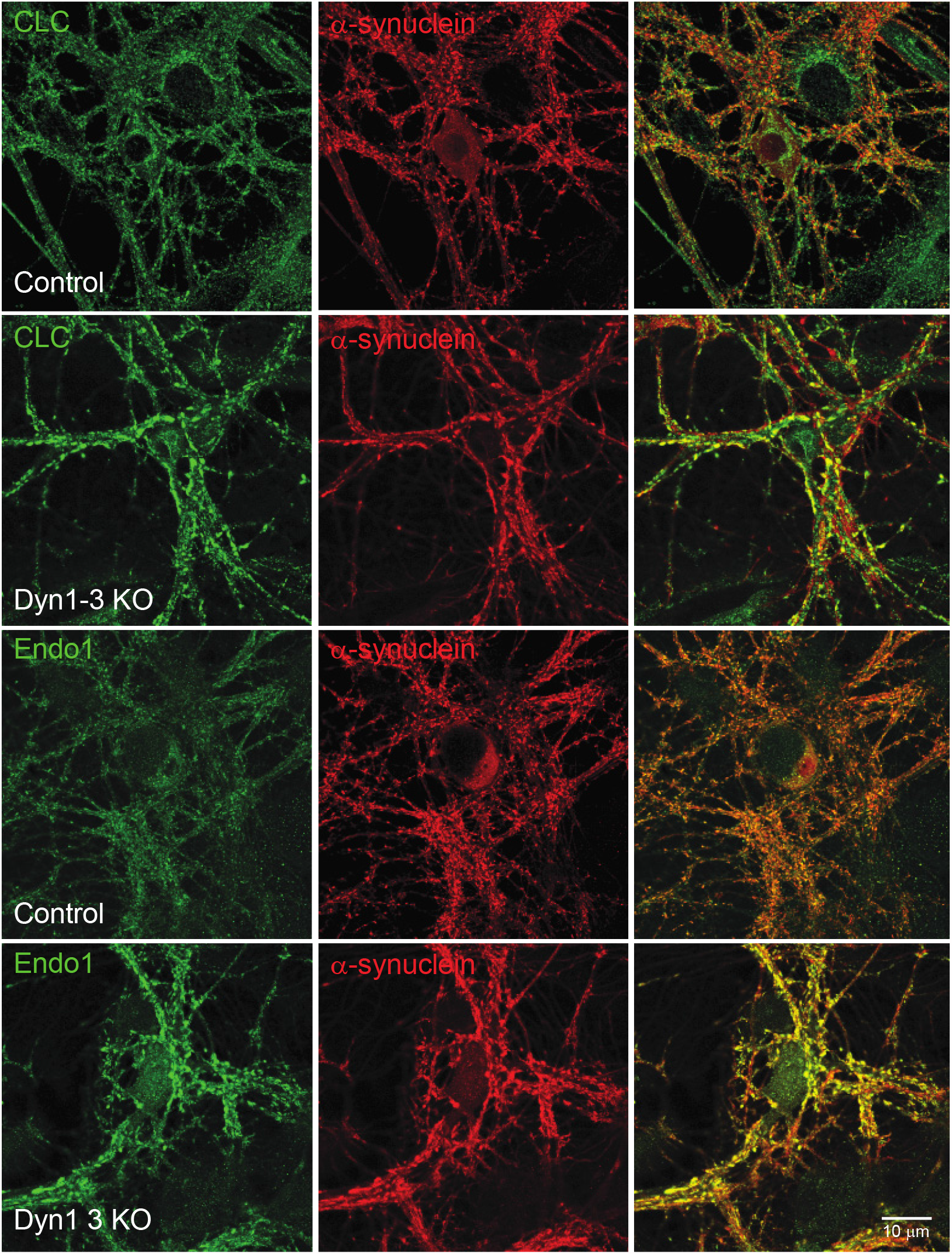
α-Synuclein is localized to endocytic clusters in Dynamin 1,3 double KO neurons. Control and Dynamin 1,3 double KO neurons were immunostained with antibodies to an endocytic protein (clathrin-CLC or Endophilin A1-Endo1) and α-synuclein. Clustering of endocytic proteins is observed in the Dynamin 1,3 double KO but not in the Control. α-Synuclein colocalizes with other trapped endocytic proteins. Scale bar =10 um.

**Supplementary Figure 2.**
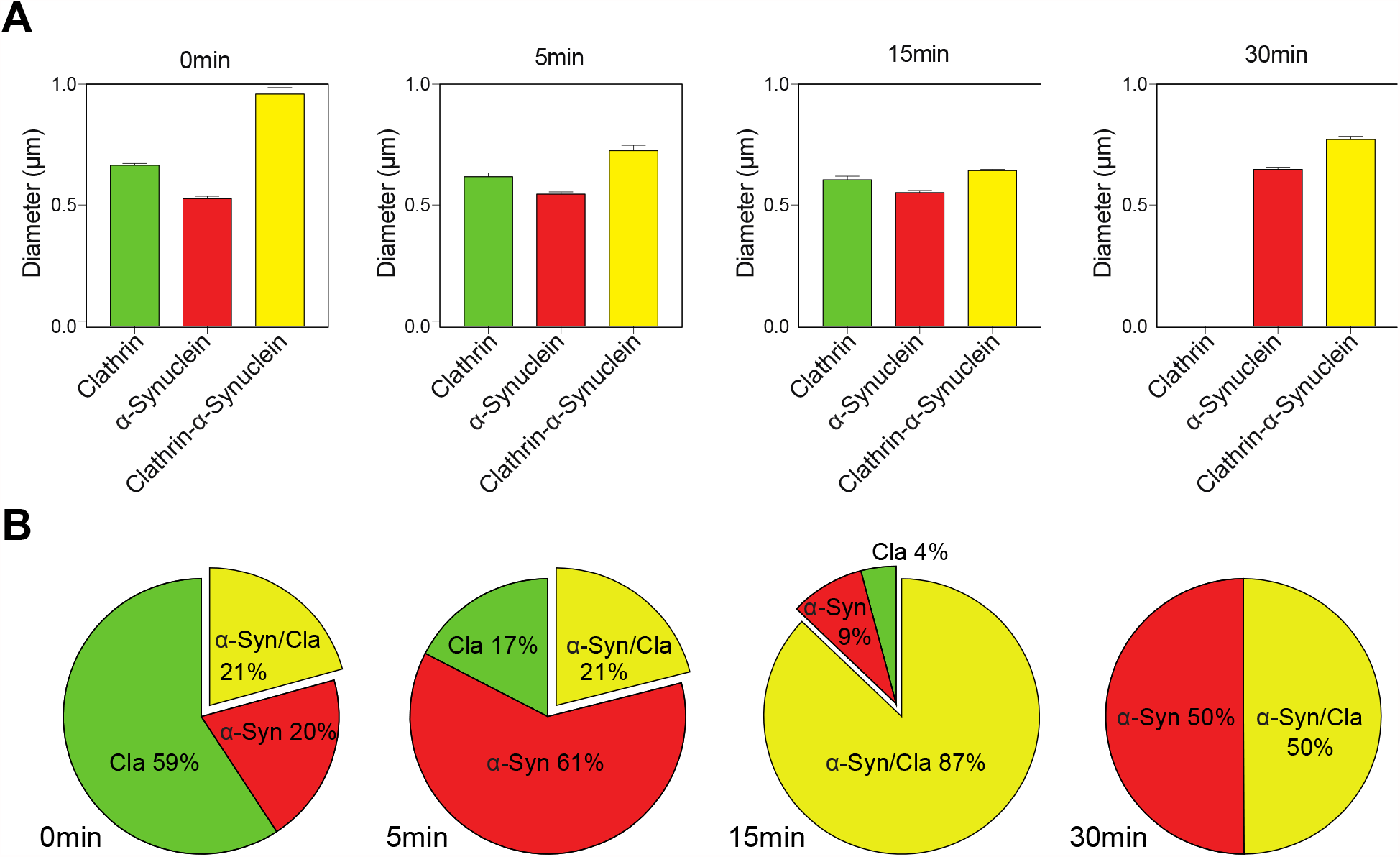
Temporal changes of size and colocalization of α-synuclein and clathrin patches. PTK2 cells were sheared and processed to detect α-synuclein and clathrin on the resulting membrane sheets. **(A)** Size comparisons of clathrin only, α-synuclein only, and clathrin+α-synuclein patches at the denoted time points. Welsh’s t-test was used to determine significance. * p<0.05; ** p<0.01 and *** p<0.0001. **(B)** Fraction of the three type of patches as a function of time.

**Supplementary Figure 3.**
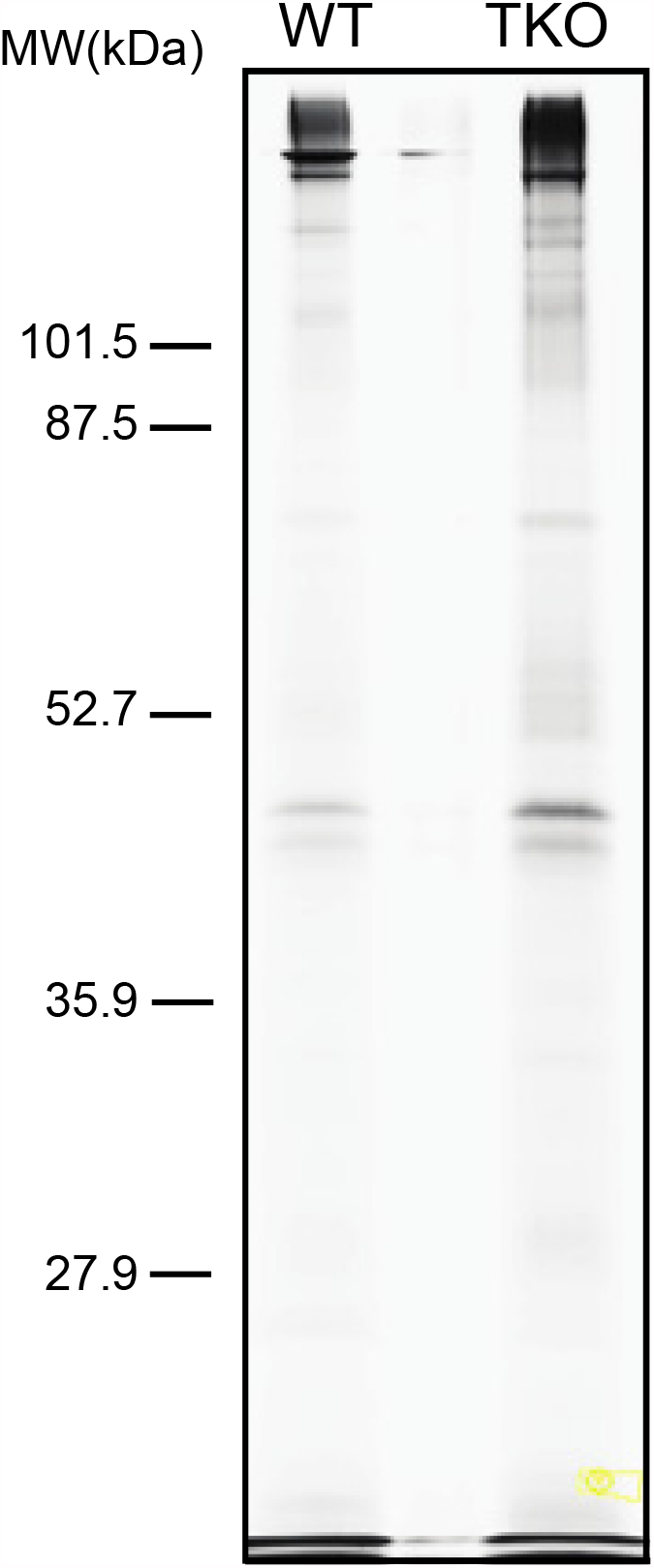
Immunoprecipitation of crosslinked α-synuclein from WT and αβγ-synuclein KO synaptosomes. WT and αβγ-synuclein KO synaptosomes were crosslinked, solubilized and immunoprecipitated with an antibody against α-synuclein. The immunoprecipitants were load in a polyacrylamide gel and stained with silver. Bands marked with boxes were cut and processed for mass spectroscopy.

## References

Antonny, B. (2011). Mechanisms of membrane curvature sensing. Annu Rev Biochem, 80, 101–123. https://doi.org/10.1146/annurev-biochem-052809-155121

Anwar, S., Peters, O., Millership, S., Ninkina, N., Doig, N., Connor-Robson, N., Threlfell, S., Kooner, G., Deacon, R. M., Bannerman, D. M., Bolam, J. P., Chandra, S. S., Cragg, S. J., Wade-Martins, R., & Buchman, V. L. (2011). Functional Alterations to the Nigrostriatal System in Mice Lacking All Three Members of the Synuclein Family. Journal of Neuroscience, 31(20), 7264–7274. https://doi.org/10.1523/jneurosci.6194-10.2011

Atias, M., Tevet, Y., Sun, J., Stavsky, A., Tal, S., Kahn, J., Roy, S., & Gitler, D. (2019). Synapsins regulate α-synuclein functions. Proceedings of the National Academy of Sciences, 116(23), 11116–11118. https://doi.org/10.1073/pnas.1903054116

Boassa, D., Berlanga, M. L., Yang, M. A., Terada, M., Hu, J., Bushong, E. A., Hwang, M., Masliah, E., George, J. M., & Ellisman, M. H. (2013). Mapping the subcellular distribution of alpha-synuclein in neurons using genetically encoded probes for correlated light and electron microscopy: implications for Parkinson’s disease pathogenesis. J Neurosci, 33(6), 2605–2615. https://doi.org/10.1523/JNEUROSCI.2898-12.2013

Brundin, P., Dave, K. D., & Kordower, J. H. (2017). Therapeutic approaches to target alpha-synuclein pathology. Exp Neurol, 298(Pt B), 225–235. https://doi.org/10.1016/j.expneurol.2017.10.003

Bucher, D., Frey, F., Sochacki, K. A., Kummer, S., Bergeest, J. P., Godinez, W. J., Krausslich, H. G., Rohr, K., Taraska, J. W., Schwarz, U. S., & Boulant, S. (2018). Clathrin-adaptor ratio and membrane tension regulate the flat-to-curved transition of the clathrin coat during endocytosis. Nat Commun, 9(1), 1109. https://doi.org/10.1038/s41467-018-03533-0

Burre, J., Sharma, M., & Sudhof, T. C. (2012). Systematic mutagenesis of alpha-synuclein reveals distinct sequence requirements for physiological and pathological activities. J Neurosci, 32(43), 15227–15242. https://doi.org/10.1523/JNEUROSCI.3545-12.2012

Burré, J., Sharma, M., & Südhof, T. C. (2014). α-Synuclein assembles into higher-order multimers upon membrane binding to promote SNARE complex formation. Proceedings of the National Academy of Sciences, 111(40), E4274–E4283. https://doi.org/10.1073/pnas.1416598111

Burre, J., Sharma, M., Tsetsenis, T., Buchman, V., Etherton, M. R., & Sudhof, T. C. (2010). Alpha-synuclein promotes SNARE-complex assembly in vivo and in vitro. Science, 329(5999), 1663–1667. https://doi.org/10.1126/science.1195227

Busch, D. J., Oliphint, P. A., Walsh, R. B., Banks, S. M., Woods, W. S., George, J. M., & Morgan, J. R. (2014). Acute increase of alpha-synuclein inhibits synaptic vesicle recycling evoked during intense stimulation. Mol Biol Cell, 25(24), 3926–3941. https://doi.org/10.1091/mbc.E14-02-0708

Chandra, S., Chen, X., Rizo, J., Jahn, R., & Sudhof, T. C. (2003). A broken alpha-helix in folded alpha-Synuclein. J Biol Chem, 278(17), 15313–15318. https://doi.org/10.1074/jbc.M213128200

Chandra, S., Gallardo, G., Fernandez-Chacon, R., Schluter, O. M., & Sudhof, T. C. (2005). Alpha-synuclein cooperates with CSPalpha in preventing neurodegeneration. Cell, 123(3), 383–396. https://doi.org/10.1016/j.cell.2005.09.028

Chen, Y., Yong, J., Martinez-Sanchez, A., Yang, Y., Wu, Y., De Camilli, P., Fernandez-Busnadiego, R., & Wu, M. (2019). Dynamic instability of clathrin assembly provides proofreading control for endocytosis. J Cell Biol, 218(10), 3200–3211. https://doi.org/10.1083/jcb.201804136

Chung, C. Y., Khurana, V., Yi, S., Sahni, N., Loh, K. H., Auluck, P. K., Baru, V., Udeshi, N. D., Freyzon, Y., Carr, S. A., Hill, D. E., Vidal, M., Ting, A. Y., & Lindquist, S. (2017). In Situ Peroxidase Labeling and Mass-Spectrometry Connects Alpha-Synuclein Directly to Endocytic Trafficking and mRNA Metabolism in Neurons. Cell Syst, 4(2), 242–250 e244. https://doi.org/10.1016/j.cels.2017.01.002

Clayton, D. F., & George, J. M. (1999). Synucleins in synaptic plasticity and neurodegenerative disorders. J Neurosci Res, 58(1), 120–129. https://www.ncbi.nlm.nih.gov/pubmed/10491577

De Camilli, P., Harris, S. M., Jr., Huttner, W. B., & Greengard, P. (1983). Synapsin I (Protein I), a nerve terminal-specific phosphoprotein. II. Its specific association with synaptic vesicles demonstrated by immunocytochemistry in agarose-embedded synaptosomes. J Cell Biol, 96(5), 1355–1373. https://doi.org/10.1083/jcb.96.5.1355

Diao, J., Burre, J., Vivona, S., Cipriano, D. J., Sharma, M., Kyoung, M., Sudhof, T. C., & Brunger, A. T. (2013). Native alpha-synuclein induces clustering of synaptic-vesicle mimics via binding to phospholipids and synaptobrevin-2/VAMP2. Elife, 2, e00592. https://doi.org/10.7554/eLife.00592

Evergren, E., Tomilin, N., Vasylieva, E., Sergeeva, V., Bloom, O., Gad, H., Capani, F., & Shupliakov, O. (2004). A pre-embedding immunogold approach for detection of synaptic endocytic proteins in situ. J Neurosci Methods, 135(1-2), 169–174. https://doi.org/10.1016/j.jneumeth.2003.12.010

Ferese, R., Modugno, N., Campopiano, R., Santilli, M., Zampatti, S., Giardina, E., Nardone, A., Postorivo, D., Fornai, F., Novelli, G., Romoli, E., Ruggieri, S., & Gambardella, S. (2015). Four Copies of SNCA Responsible for Autosomal Dominant Parkinson’s Disease in Two Italian Siblings. Parkinsons Dis, 2015, 546462. https://doi.org/10.1155/2015/546462

Ferguson, S. M., Brasnjo, G., Hayashi, M., Wolfel, M., Collesi, C., Giovedi, S., Raimondi, A., Gong, L. W., Ariel, P., Paradise, S., O’Toole, E., Flavell, R., Cremona, O., Miesenbock, G., Ryan, T. A., & De Camilli, P. (2007). A selective activity-dependent requirement for dynamin 1 in synaptic vesicle endocytosis. Science, 316(5824), 570–574. https://doi.org/10.1126/science.1140621

Ferreon, A. C., Gambin, Y., Lemke, E. A., & Deniz, A. A. (2009). Interplay of alpha-synuclein binding and conformational switching probed by single-molecule fluorescence. Proc Natl Acad Sci U S A, 106(14), 5645–5650. https://doi.org/10.1073/pnas.0809232106

Ford, M. G., Pearse, B. M., Higgins, M. K., Vallis, Y., Owen, D. J., Gibson, A., Hopkins, C. R., Evans, P. R., & McMahon, H. T. (2001). Simultaneous binding of PtdIns(4,5)P2 and clathrin by AP180 in the nucleation of clathrin lattices on membranes. Science, 291(5506), 1051–1055. https://doi.org/10.1126/science.291.5506.1051

Fortin, D. L., Nemani, V. M., Voglmaier, S. M., Anthony, M. D., Ryan, T. A., & Edwards, R. H. (2005). Neural activity controls the synaptic accumulation of alpha-synuclein. J Neurosci, 25(47), 10913–10921. https://doi.org/10.1523/JNEUROSCI.2922-05.2005

Greten-Harrison, B., Polydoro, M., Morimoto-Tomita, M., Diao, L., Williams, A. M., Nie, E. H., Makani, S., Tian, N., Castillo, P. E., Buchman, V. L., & Chandra, S. S. (2010). alphabetagamma-Synuclein triple knockout mice reveal age-dependent neuronal dysfunction. Proc Natl Acad Sci U S A, 107(45), 19573–19578. https://doi.org/10.1073/pnas.1005005107

Hosaka, M., Hammer, R. E., & Sudhof, T. C. (1999). A phospho-switch controls the dynamic association of synapsins with synaptic vesicles. Neuron, 24(2), 377–387. https://www.ncbi.nlm.nih.gov/pubmed/10571231

Ibanez, P., Bonnet, A. M., Debarges, B., Lohmann, E., Tison, F., Pollak, P., Agid, Y., Durr, A., & Brice, A. (2004). Causal relation between alpha-synuclein gene duplication and familial Parkinson’s disease. Lancet, 364(9440), 1169–1171. https://doi.org/10.1016/S0140-6736(04)17104-3

Jao CC, Hegde BG, Chen J, Haworth IS, Langen R. (2008). Structure of membrane-bound alpha-synuclein from site-directed spin labeling and computational refinement. Proc Natl Acad Sci U S A, 105(50):19666–71. doi: 10.1073/pnas.0807826105.

Kaur, U., & Lee, J. C. (2020). Unroofing site-specific alpha-synuclein-lipid interactions at the plasma membrane. Proc Natl Acad Sci U S A, 117(32), 18977–18983. https://doi.org/10.1073/pnas.2006291117

Koo, S. J., Kochlamazashvili, G., Rost, B., Puchkov, D., Gimber, N., Lehmann, M., Tadeus, G., Schmoranzer, J., Rosenmund, C., Haucke, V., & Maritzen, T. (2015). Vesicular Synaptobrevin/VAMP2 Levels Guarded by AP180 Control Efficient Neurotransmission. Neuron, 88(2), 330–344. https://doi.org/10.1016/j.neuron.2015.08.034

Koo, S. J., Markovic, S., Puchkov, D., Mahrenholz, C. C., Beceren-Braun, F., Maritzen, T., Dernedde, J., Volkmer, R., Oschkinat, H., & Haucke, V. (2011). SNARE motif-mediated sorting of synaptobrevin by the endocytic adaptors clathrin assembly lymphoid myeloid leukemia (CALM) and AP180 at synapses. Proc Natl Acad Sci U S A, 108(33), 13540–13545. https://doi.org/10.1073/pnas.1107067108

Koo, S. J., Puchkov, D., & Haucke, V. (2011). AP180 and CALM: Dedicated endocytic adaptors for the retrieval of synaptobrevin 2 at synapses. Cell Logist, 1(4), 168–172. https://doi.org/10.4161/cl.1.4.18897

Kruger, R., Kuhn, W., Muller, T., Woitalla, D., Graeber, M., Kosel, S., Przuntek, H., Epplen, J. T., Schols, L., & Riess, O. (1998). Ala30Pro mutation in the gene encoding alpha-synuclein in Parkinson’s disease. Nat Genet, 18(2), 106–108. https://doi.org/10.1038/ng0298-106

Logan, T., Bendor, J., Toupin, C., Thorn, K., & Edwards, R. H. (2017). alpha-Synuclein promotes dilation of the exocytotic fusion pore. Nat Neurosci, 20(5), 681–689. https://doi.org/10.1038/nn.4529

Lokappa, S. B., & Ulmer, T. S. (2011). Alpha-synuclein populates both elongated and broken helix states on small unilamellar vesicles. J Biol Chem, 286(24), 21450–21457. https://doi.org/10.1074/jbc.M111.224055

Maroteaux, L., Campanelli, J., & Scheller, R. (1988). Synuclein: a neuron-specific protein localized to the nucleus and presynaptic nerve terminal. The Journal of Neuroscience, 8(8), 2804–2815. https://doi.org/10.1523/jneurosci.08-08-02804.1988

Medeiros, A. T., Bubacco, L., & Morgan, J. R. (2018). Impacts of increased α-synuclein on clathrin-mediated endocytosis at synapses: implications for neurodegenerative diseases. Neural Regen Res, 13(4), 647–648. https://doi.org/10.4103/1673-5374.230289

Middleton, E. R., & Rhoades, E. (2010). Effects of curvature and composition on alpha-synuclein binding to lipid vesicles. Biophys J, 99(7), 2279–2288. https://doi.org/10.1016/j.bpj.2010.07.056

Moshkanbaryans, L., Chan, L. S., & Graham, M. E. (2014). The Biochemical Properties and Functions of CALM and AP180 in Clathrin Mediated Endocytosis. Membranes (Basel), 4(3), 388–413. https://doi.org/10.3390/membranes4030388

Pechstein, A., Bacetic, J., Vahedi-Faridi, A., Gromova, K., Sundborger, A., Tomlin, N., Krainer, G., Vorontsova, O., Schafer, J. G., Owe, S. G., Cousin, M. A., Saenger, W., Shupliakov, O., & Haucke, V. (2010). Regulation of synaptic vesicle recycling by complex formation between intersectin 1 and the clathrin adaptor complex AP2. Proc Natl Acad Sci U S A, 107(9), 4206–4211. https://doi.org/10.1073/pnas.0911073107

Polymeropoulos, M. H. (1997). Mutation in the -Synuclein Gene Identified in Families with Parkinson’s Disease. Science, 276(5321), 2045–2047. https://doi.org/10.1126/science.276.5321.2045

Raimondi, A., Ferguson, S. M., Lou, X., Armbruster, M., Paradise, S., Giovedi, S., Messa, M., Kono, N., Takasaki, J., Cappello, V., O’Toole, E., Ryan, T. A., & De Camilli, P. (2011). Overlapping role of dynamin isoforms in synaptic vesicle endocytosis. Neuron, 70(6), 1100–1114. https://doi.org/10.1016/j.neuron.2011.04.031

Rhoades, E., Ramlall, T. F., Webb, W. W., & Eliezer, D. (2006). Quantification of alpha-synuclein binding to lipid vesicles using fluorescence correlation spectroscopy. Biophys J, 90(12), 4692–4700. https://doi.org/10.1529/biophysj.105.079251

Ringstad, N., Gad, H., Low, P., Di Paolo, G., Brodin, L., Shupliakov, O., & De Camilli, P. (1999). Endophilin/SH3p4 is required for the transition from early to late stages in clathrin-mediated synaptic vesicle endocytosis. Neuron, 24(1), 143–154. https://www.ncbi.nlm.nih.gov/pubmed/10677033

Saffarian, S., Cocucci, E., & Kirchhausen, T. (2009). Distinct dynamics of endocytic clathrin-coated pits and coated plaques. PLoS Biol, 7(9), e1000191. https://doi.org/10.1371/journal.pbio.1000191

Satake, W., Nakabayashi, Y., Mizuta, I., Hirota, Y., Ito, C., Kubo, M., Kawaguchi, T., Tsunoda, T., Watanabe, M., Takeda, A., Tomiyama, H., Nakashima, K., Hasegawa, K., Obata, F., Yoshikawa, T., Kawakami, H., Sakoda, S., Yamamoto, M., Hattori, N., Murata, M., Nakamura, Y., & Toda, T. (2009). Genome-wide association study identifies common variants at four loci as genetic risk factors for Parkinson’s disease. Nat Genet, 41(12), 1303–1307. https://doi.org/10.1038/ng.485

Schechter, M., Atias, M., Abd Elhadi, S., Davidi, D., Gitler, D., & Sharon, R. (2020). alpha-Synuclein facilitates endocytosis by elevating the steady-state levels of phosphatidylinositol 4,5-bisphosphate. J Biol Chem, 295(52), 18076–18090. https://doi.org/10.1074/jbc.RA120.015319

Simon-Sanchez, J., Schulte, C., Bras, J. M., Sharma, M., Gibbs, J. R., Berg, D., Paisan-Ruiz, C., Lichtner, P., Scholz, S. W., Hernandez, D. G., Kruger, R., Federoff, M., Klein, C., Goate, A., Perlmutter, J., Bonin, M., Nalls, M. A., Illig, T., Gieger, C., Houlden, H., Steffens, M., Okun, M. S., Racette, B. A., Cookson, M. R., Foote, K. D., Fernandez, H. H., Traynor, B. J., Schreiber, S., Arepalli, S., Zonozi, R., Gwinn, K., van der Brug, M., Lopez, G., Chanock, S. J., Schatzkin, A., Park, Y., Hollenbeck, A., Gao, J., Huang, X., Wood, N. W., Lorenz, D., Deuschl, G., Chen, H., Riess, O., Hardy, J. A., Singleton, A. B., & Gasser, T. (2009). Genome-wide association study reveals genetic risk underlying Parkinson’s disease. Nat Genet, 41(12), 1308–1312. https://doi.org/10.1038/ng.487

Singleton, A. B., Farrer, M., Johnson, J., Singleton, A., Hague, S., Kachergus, J., Hulihan, M., Peuralinna, T., Dutra, A., Nussbaum, R., Lincoln, S., Crawley, A., Hanson, M., Maraganore, D., Adler, C., Cookson, M. R., Muenter, M., Baptista, M., Miller, D., Blancato, J., Hardy, J., & Gwinn-Hardy, K. (2003). alpha-Synuclein locus triplication causes Parkinson’s disease. Science, 302(5646), 841. https://doi.org/10.1126/science.1090278

Sochacki, K. A., Dickey, A. M., Strub, M. P., & Taraska, J. W. (2017). Endocytic proteins are partitioned at the edge of the clathrin lattice in mammalian cells. Nat Cell Biol, 19(4), 352–361. https://doi.org/10.1038/ncb3498

Sochacki, K. A., & Taraska, J. W. (2017). Correlative Fluorescence Super-Resolution Localization Microscopy and Platinum Replica EM on Unroofed Cells. Methods Mol Biol, 1663, 219–230. https://doi.org/10.1007/978-1-4939-7265-4_18

Spillantini, M. G., Schmidt, M. L., Lee, V. M. Y., Trojanowski, J. Q., Jakes, R., & Goedert, M. (1997). α-Synuclein in Lewy bodies. Nature, 388(6645), 839–840. https://doi.org/10.1038/42166

Sun, J., Wang, L., Bao, H., Premi, S., Das, U., Chapman, E. R., & Roy, S. (2019). Functional cooperation of alpha-synuclein and VAMP2 in synaptic vesicle recycling. Proc Natl Acad Sci U S A, 116(23), 11113–11115. https://doi.org/10.1073/pnas.1903049116

Taguchi, Y. V., Liu, J., Ruan, J., Pacheco, J., Zhang, X., Abbasi, J., Keutzer, J., Mistry, P. K., & Chandra, S. S. (2017). Glucosylsphingosine Promotes alpha-Synuclein Pathology in Mutant GBA-Associated Parkinson’s Disease. J Neurosci, 37(40), 9617–9631. https://doi.org/10.1523/JNEUROSCI.1525-17.2017

Takamori, S., Holt, M., Stenius, K., Lemke, E. A., Gronborg, M., Riedel, D., Urlaub, H., Schenck, S., Brugger, B., Ringler, P., Muller, S. A., Rammner, B., Grater, F., Hub, J. S., De Groot, B. L., Mieskes, G., Moriyama, Y., Klingauf, J., Grubmuller, H., Heuser, J., Wieland, F., & Jahn, R. (2006). Molecular anatomy of a trafficking organelle. Cell, 127(4), 831–846. https://doi.org/10.1016/j.cell.2006.10.030

Tonges, L., & Zella, M. A. S. (2019). Antibody-based immunotherapies for Parkinsonian syndromes. Neural Regen Res, 14(11), 1903–1904. https://doi.org/10.4103/1673-5374.259613

Vargas, K. J., Makani, S., Davis, T., Westphal, C. H., Castillo, P. E., & Chandra, S. S. (2014). Synucleins regulate the kinetics of synaptic vesicle endocytosis. J Neurosci, 34(28), 9364–9376. https://doi.org/10.1523/JNEUROSCI.4787-13.2014

Vargas, K. J., Schrod, N., Davis, T., Fernandez-Busnadiego, R., Taguchi, Y. V., Laugks, U., Lucic, V., & Chandra, S. S. (2017). Synucleins Have Multiple Effects on Presynaptic Architecture. Cell Rep, 18(1), 161–173. https://doi.org/10.1016/j.celrep.2016.12.023

Wang, L., Das, U., Scott, D. A., Tang, Y., McLean, P. J., & Roy, S. (2014). alpha-synuclein multimers cluster synaptic vesicles and attenuate recycling. Curr Biol, 24(19), 2319–2326. https://doi.org/10.1016/j.cub.2014.08.027

Westphal, C. H., & Chandra, S. S. (2013). Monomeric synucleins generate membrane curvature. J Biol Chem, 288(3), 1829–1840. https://doi.org/10.1074/jbc.M112.418871

Wu, M., Huang, B., Graham, M., Raimondi, A., Heuser, J. E., Zhuang, X., & De Camilli, P. (2010). Coupling between clathrin-dependent endocytic budding and F-BAR-dependent tubulation in a cell-free system. Nat Cell Biol, 12(9), 902–908. https://doi.org/10.1038/ncb2094

Xu, J., Wu, X. S., Sheng, J., Zhang, Z., Yue, H. Y., Sun, L., Sgobio, C., Lin, X., Peng, S., Jin, Y., Gan, L., Cai, H., & Wu, L. G. (2016). α-Synuclein Mutation Inhibits Endocytosis at Mammalian Central Nerve Terminals. J Neurosci, 36(16), 4408–4414. https://doi.org/10.1523/jneurosci.3627-15.2016

Zarranz, J. J., Alegre, J., Gomez-Esteban, J. C., Lezcano, E., Ros, R., Ampuero, I., Vidal, L., Hoenicka, J., Rodriguez, O., Atares, B., Llorens, V., Gomez Tortosa, E., del Ser, T., Munoz, D. G., & de Yebenes, J. G. (2004). The new mutation, E46K, of alpha-synuclein causes Parkinson and Lewy body dementia. Ann Neurol, 55(2), 164–173. https://doi.org/10.1002/ana.10795

